# Chromatin access regulates the formation of Müller glia-derived progenitor cells in the retina

**DOI:** 10.1101/2022.06.23.497349

**Authors:** Warren A. Campbell, Heithem M. El-Hodiri, Diego Torres, Evan C. Hawthorn, Lisa E. Kelly, Leo Volkov, David Akanonu, Andy J. Fischer

## Abstract

Chromatin access and epigenetic control over gene expression play important roles in regulating developmental processes. However, little is known about how chromatin access and epigenetic gene silencing influence mature glial cells and retinal regeneration. Herein we investigate the expression and functions of S-Adenosylhomocysteine Hydrolase (SAHH; *AHCY*) and Histone Methyltransferases (HMTs) during the formation of Müller glia-derived progenitor cells (MGPCs) in the chick and mouse retinas. In chick, *AHCY, AHCYL1, AHCYL2* and many different HMTs are dynamically expressed by MG and MGPCs in damaged retinas. Inhibition of SAHH reduced levels of H3K27me3 and potently blocks the formation of proliferating MGPCs. By using a combination of single cell RNA-seq and single cell ATAC-seq, we find significant changes in gene expression and chromatin access in MG with SAHH inhibition and NMDA-treatment; many of these genes are associated with glial and neuronal differentiation. A strong correlation across gene expression, chromatin access, and transcription factor motif access in MG was observed for transcription factors known to covey glial identity and promote retinal development. By comparison, in the mouse retina, inhibition of SAHH has no influence on the formation of neuron-like cells from *Ascl1*-overexpressing MG. We conclude that in the chick, but not the mouse, the activity of SAHH and HMTs are required for the reprogramming of MG into MGPCs by regulating chromatin access to transcription factors associated with glial differentiation and retinal development.

## Introduction

Müller glia (MG) are the predominant type of support cell in the retina and are common to all vertebrate classes. MG perform many important functions including structural support, synaptic support, osmotic homeostasis, and metabolic support to retinal neurons (Bringmann et al., 2006; Bringmann et al., 2009). However, MG also have the extraordinary capability to become progenitor cells that regenerate retinal neurons (Bernardos et al., 2007; Fausett et al., 2008; Fischer and Reh, 2001; Karl et al., 2008; Ooto et al., 2004). The transition of MG to Müller glia-derived progenitor cells (MGPCs) involves downregulation of glial genes, upregulation of progenitor-associated factors and proliferation followed by neuronal differentiation (Fischer and Bongini, 2010; Gallina et al., 2014a; Todd and Reh, 2021; Wilken and Reh, 2016). The process of retinal regeneration is highly efficient in fish, whereas this process is sadly ineffective in mammals. In the mouse retina, significant stimulation is required to induce the formation of MGPCs. For example, the forced expression of the transcription factor *Ascl1*, in combination with neuronal damage and inhibition of Histone Deacetylases (HDACs), stimulates reprogramming of MG into functional, light-responsive neurons (Jorstad et al., 2017; Pollak et al., 2013; Ueki et al., 2015). The differentiation of neurons from *Ascl1* overexpressing MG can be enhanced by ablating reactive microglia (Todd et al., 2020); by inhibiting Jak/Stat-signaling (Jorstad et al., 2020), NFkB-signaling, TGFβ/Smad3-signaling or ID transcription factors (Palazzo et al., 2022); or by combined over-expression of *Ascl1* with *Atoh1* (Todd et al., 2021). By comparison, the process of retinal regeneration in birds is limited; numerous proliferating MGPCs are formed, but few of the progeny differentiate into neurons (reviewed by (Gallina et al., 2014a). Accordingly, the chick retina represents a valuable model in which to study factors that activate or suppress the formation of neurogenic MGPCs. Although many different factors have been identified that influence the reprogramming of MG into proliferating, neurogenic progenitor cells, little is known about how epigenetic mechanisms regulate chromatin access to influence the process of reprogramming.

The purpose of this study was to investigate how the activity of S-Adenosylhomocysteine Hydrolase (SAHH) and downstream methyltransferases regulate chromatin access and the reprogramming of MG into proliferating MGPCs. The *AHCY* gene codes for S-Adenosylhomocysteine Hydrolase (SAHH), *AHCYL1* codes for SAHH2 protein, and *AHCYL2* codes for SAHH3 protein. SAHH enzymes catalyze the hydrolysis of S-adenosylhomocysteine (SAH) to adenosine and L-homocysteine. Inhibition of SAHH enzymes causes the accumulation of intracellular SAH which potently inhibits methyl transfer reactions (Hershfield et al., 1979), namely the activity Histone Methyltransferases (HMTs) (reviewed by (Vizán et al., 2021). Thus, inhibition of SAHH activity is an effective method to broadly disrupt HMTs and influence chromatin access (Miranda et al., 2009). HMTs catalyze the transfer of methyl groups to basic amino acid residues of histones, predominantly histone H3 and H4 (Wood and Shilatifard, 2004). There are 2 major types of HMTs: one type selectively methylates arginine residues; the other type methylates lysine residues, which includes HMTs such as SET domain-containing (such as SUV39H and Enhancer of Zeste H2) or non-SET domain-containing enzymes (Feng et al., 2002; Ng et al., 2002; Sawan and Herceg, 2010). Histone methylation is an important modification of chromatin that epigenetically regulates levels of gene expression, genomic stability, stem cell function, cell lineage development, cellular differentiation and cell division.

Currently, little is known about the changes in chromatin access that occur in MG during reprogramming into MGPCs. Further, nothing is known about how changes in chromatin access that are regulated by SAHH and HMTs influence mature MG and the formation of MGPCs. Accordingly, we investigated whether changes in chromatin access occur in MG during the process of reprogramming and whether inhibition SAHH influences glial phenotype, de-differentiation and the formation of MGPCs *in vivo*.

## Methods and Materials

### Animals

The animals approved for use in these experiments was in accordance with the guidelines established by the National Institutes of Health and IACUC at The Ohio State University. Newly hatched P0 wildtype leghorn chicks (*Gallus gallus domesticus*) were obtained from Meyer Hatchery (Polk, Ohio). Post-hatch chicks were maintained in a regular diurnal cycle of 12 hours light, 12 hours dark (lights on at 8:00 AM). Chicks were housed in stainless-steel brooders at 25°C and received water and Purina^tm^ chick starter *ad libitum*.

Mice were kept on a cycle of 12 hours light, 12 hours dark (lights on at 8:00 AM). The use of Ascl1 over-expressing mice (*Glast-CreER:LNL-tTA:tetO-mAscl1-ires-GFP)* was as previously described (Jorstad et al., 2017; Palazzo et al., 2022).

### Intraocular injections

Chicks were anesthetized with 2.5% isoflurane mixed with oxygen from a non-rebreathing vaporizer. The technical procedures for intraocular injections were performed as previously described (Fischer et al., 1998). With all injection paradigms, both pharmacological and vehicle treatments were administered to the right and left eye respectively. Compounds were injected in 20 µl sterile saline with 0.05 mg/ml bovine serum albumin added as a carrier. Compounds included: NMDA (38.5nmol or 154 µg/dose; Sigma-Aldrich), FGF2 (250 ng/dose; R&D systems), and DZNexp4 (10 µg/dose in DMSO; Tocris). 5-Ethynyl-2’-deoxyuridine (EdU; 2.3 µg/dose) was intravitreally injected to label the nuclei of proliferating cells. Injection paradigms are included in each figure.

Mice were anesthetized via inhalation of 2.5% isoflurane in oxygen and intraocular injections performed as described previously (Palazzo et al., 2022; Todd et al., 2015; Todd et al., 2019). Right eyes of mice were injected with the experimental compound and the contra-lateral left eyes were injected with a control vehicle. Compounds used in these studies included N-methyl-D-aspartate (NMDA; Sigma; 1.5 µl at concentration of 100mM in PBS), Trichostatin A (TSA; Sigma; 1 µg/dose in DMSO), and 3-Deazaneplanocin A hydrochloride (DZN; 1 µg/dose in DMSO; Tocris). Tamoxifen (Sigma; 1.5 mg/100 µl corn oil per dose) was applied as 4 consecutive daily intraperitoneal injections.

### Single Cell RNA and ATAC sequencing of retinas

Retinas were obtained from adult mice and post-hatch chicks. For chicks 2 retinas were pooled for each library/experimental condition. For mice, 3 retinas were pooled for each library/experimental condition. Isolated retinas were dissociated in a 0.25% papain solution in Hank’s balanced salt solution (HBSS), pH = 7.4, for 30 minutes, and suspensions were frequently triturated. The dissociated cells were passed through a sterile 70µm filter to remove large particulate debris. Dissociated cells were assessed for viability (Countess II; Invitrogen) and cell-density diluted to 700 cell/µl. Each single cell cDNA library was prepared for a target of 10,000 cells per sample. The cell suspension and Chromium Single Cell 3’ V2 or V3 reagents (10X Genomics) were loaded onto chips to capture individual cells with individual gel beads in emulsion (GEMs) using 10X Chromium Controller. cDNA and library amplification for an optimal signal was 12 and 10 cycles respectively. Sequencing was conducted on Illumina HiSeq2500 (Genomics Resource Core Facility, John’s Hopkins University) or HiSeq4000/Novaseq (Novogene) with 2x ≤ 100 bp paired-end reads. Fasta sequence files were de-multiplexed, aligned, and annotated using the chick ENSMBL database (Chick-5.0, Ensembl release 94) and Cell Ranger software. Gene expression was counted using unique molecular identifier bar codes, and gene-cell matrices were constructed. Using Seurat and Signac toolkits, Uniform Manifold Approximation and Projection for Dimension Reduction (UMAP) plots were generated from aggregates of multiple scRNA-seq libraries (Butler et al., 2018; Satija et al., 2015). Seurat was used to construct violin/scatter plots. Significance of difference in violin/scatter plots was determined using a Wilcoxon Rank Sum test with Bonferroni correction. Monocle was used to construct unbiased pseudo-time trajectories and scatter plotters for MG and MGPCs across pseudotime (Qiu et al., 2017a; Qiu et al., 2017b; Trapnell et al., 2012). SingleCellSignalR was used to assess potential ligand-receptor interactions between cells within scRNA-seq datasets (Cabello-Aguilar et al., 2020). To perform Gene Ontology (GO) enrichment, analyses lists of DEGs were uploaded to ShinyGo v0.66 (http://bioinformatics.sdstate.edu/go/).

Genes that were used to identify different types of retinal cells included the following: (i) Müller glia: *GLUL, VIM, SCL1A3, RLBP1*; (ii) MGPCs: *PCNA, CDK1, TOP2A, ASCL1*; (iii) microglia: *C1QA, C1QB, CCL4, CSF1R, TMEM22*; (iv) ganglion cells: *THY1, POU4F2, RBPMS2, NEFL, NEFM*; (v) amacrine cells: *GAD67, CALB2, TFAP2A*; (vi) horizontal cells: *PROX1, CALB2, NTRK1*; (vii) bipolar cells: *VSX1, OTX2, GRIK1, GABRA1*; (viii) cone photoreceptors: *CALB1, GNAT2, OPN1LW*; and (ix) rod photoreceptors: *RHO, NR2E3, ARR3.* The MG have an over-abundant representation in the chick scRNA-seq databases. This likely resulted from a fortuitous capture-bias and/or tolerance of the MG to the dissociation process.

For scATAC-seq, FASTQ files were processed using CellRanger ATAC 2.1.0, annotated using the chick genome (Gallus_gallus-5.0, Ensembl release 84, assembly GRC6a v105), and then aggregated using CellRanger ATAC aggr to create a common peak set. CellRanger ATAC aggr output was processed and analyzed using Signac (develop version 1.6.0.9015, github.com/timoast/signac) (Stuart et al., 2021) as described elsewhere (https://satijalab.org/signac/articles/mouse_brain_vignette.html). Annotations were added as an AnnotationHub (Bioconductor.org) object generated from the Ensembl chicken genome (see above). Motif analysis was performed using Signac, the JASPAR 2020 vertebrate database (Fornes et al., 2020), and TFBSTools (Tan and Lenhard, 2016) as described in: https://satijalab.org/signac/articles/motif_vignette.html. Motif scores were generated by analysis of the Signac object using chromVAR (Schep et al., 2017). For Signac and chromVAR motif analysis, the genome used was a FaFile generated using Rsamtools (Bioconductor.org) and the necessary chicken FASTA and index files from Ensembl. For scATAC-seq analysis, including motif enrichment and motif scoring, all reference or annotation genomes were in NCBI/Ensembl format.

### Fixation, sectioning and immunocytochemistry

Retinal tissue samples were formaldehyde fixed, sectioned, and labeled via immunohistochemistry as described previously (Fischer et al., 2008a; Fischer et al., 2009c). Antibody dilutions and commercial sources used in this study are described in Table 1. Observed labeling was not due to off-target labeling of secondary antibodies or tissue autofluorescence because sections incubated with secondary antibodies alone were devoid of fluorescence. Patterns of immunolabeling for all antibodies precisely match patterns of mRNA expression in scRNA-libraries of control and damaged retinas, similar previous descriptions (Campbell et al., 2019; Campbell et al., 2021b; Campbell et al., 2021c; Todd et al., 2019). Secondary antibodies utilized include donkey-anti-goat-Alexa488/568, goat-anti-rabbit-Alexa488/568/647, goat-anti-mouse-Alexa488/568/647, goat-anti-rat-Alexa488 (Life Technologies) diluted to 1:1000 in PBS and 0.2% Triton X-100.

**Table 1.** List of antibodies, working dilution, host, clone/catalog number and source.

| Antibody | Dilution | Host | Clone/Catalog number | Source |
| --- | --- | --- | --- | --- |
| Sox2 | 1:1000 | Goat | KOY0418121 | R&D Systems |
| Sox9 | 1:2000 | Rabbit | AB5535 | Millipore |
| CD45 | 1:300 | Mouse | HIS-C7 | Cedi Diagnostic |
| PCNA | 1:1000 | Mouse | M0879 | Dako<br>Immunochemicals |
| Phospho-histone H3 | 1:600 | Rabbit |  |  |
| Neurofilament | 1:50 | Mouse | RT97 | Developmental<br>Studies Hybridoma<br>Bank (DSHB) |
| H3K37me3 |  | Mouse |  |  |
| Nkx2.2 | 1:100 | Mouse | 74.5A5 | DSHB |
| cFos |  |  |  |  |
| Glutamine synthetase | 1:1000 | Mouse | 610517 | BD Biosciences |
| pS6 | 1:750 | Rabbit | #2215; Ser240/244 | Cell Signaling |
| pStat3 | 1:100 | Rabbit | 9131 | Cell Signaling |
| pSmad1/5/8 | 1:200 | Rabbit | D5B10 | Cell Signaling |
| Anti-goat IgG Alex488 | 1:1000 | Donkey | A3214 | Life Technologies |
| Anti-goat IgG Alex568 | 1:1000 | Donkey | A-11057 | Life Technologies |
| Anti-rabbit IgG Alex488 | 1:1000 | Goat | A32731 | Life Technologies |
| Anti-rabbit IgG Alex568 | 1:1000 | Goat | A-11036 | Life Technologies |
| Anti-rabbit IgG Alex647 | 1:1000 | Goat | A32733 | Life Technologies |
| Anti-mouse IgG Alex488 | 1:1000 | Goat | A32723 | Life Technologies |
| Anti-mouse IgG Alex568 | 1:1000 | Goat | A-11004 | Life Technologies |
| Anti-mouse IgG Alex647 | 1:1000 | Goat | A32728 | Life Technologies |
| Anti-Rat IgG Alexa488 | 1:1000 | Goat | A48262 | Life Technologies |

### Labeling for EdU

For the detection of nuclei that incorporated EdU, immunolabeled sections were fixed in 4% formaldehyde in 0.1M PBS pH 7.4 for 5 minutes at room temperature. Samples were washed for 5 minutes with PBS, permeabilized with 0.5% Triton X-100 in PBS for 1 minute at room temperature and washed twice for 5 minutes in PBS. Sections were incubated for 30 minutes at room temperature in a buffer consisting of 100 mM Tris, 8 mM CuSO_4_, and 100 mM ascorbic acid in dH_2_O. The Alexa Fluor 568 Azide (Thermo Fisher Scientific) was added to the buffer at a 1:500 dilution.

### Terminal deoxynucleotidyl transferase dUTP nick end labeling (TUNEL)

The TUNEL assay was implemented to identify dying cells by imaging fluorescent labeling of double stranded DNA breaks in nuclei. The *In Situ* Cell Death Kit (TMR red; Roche Applied Science) was applied to fixed retinal sections as per the manufacturer’s instructions.

### Photography, measurements, cell counts and statistics

Microscopy images of retinal sections were captured with the Leica DM5000B microscope with epifluorescence and the Leica DC500 digital camera. High resolution confocal images were obtained with a Leica SP8 available in the Department of Neuroscience Imaging Facility at the Ohio State University. Representative images are modified to optimize color, brightness and contrast using Adobe Photoshop. For quantification of numbers of EdU^+^ cells, a fixed region of retina was counted and average numbers of Sox2+/CD45+ and EdU co-labeled cells. The retinal region selected cell counts was standardized between treatment and control groups to reduce variability and improve reproducibility.

Similar to previous reports (Fischer et al., 2009a; Fischer et al., 2009b; Ghai et al., 2009), immunofluorescence was quantified by using Image J (NIH). Identical illumination, microscope, and camera settings were used to obtain images for quantification. Measurement for immunofluorescence were made from single optical confocal sections. Measurements of pS6, pStat3 and pSmad1/5/8 immunofluorescence were made for a fixed, cropped area (25,000 µm^2^) of INL, OPL and ONL. Activation of cell signaling through mTor (pS6), Jak/Stat (pStat3), and BMP/SMAD (pSmad1/5/8) in outer layers of NMDA-damaged retinas is known to manifest exclusively in MG (Todd et al., 2016a; Todd et al., 2017; Zelinka et al., 2016). Measurements were made for regions containing pixels with intensity values of 70 or greater (0 = black and 255 = saturated). The cropped areas contain between 80 and 140 MG or MGPCs; numbers of cells vary depending on treatments that influence the proliferation of MGPCs. The intensity sum was calculated as the total of pixel values for all pixels within threshold regions. GraphPad Prism 6 was used for statistical analyses.

We performed a Levene’s test to determine whether data from control and treatment groups had equal variance. For treatment groups where the Levene’s test indicated unequal variance, we performed a Mann Whitney U test (Wilcoxon Rank Sum Test). For statistical evaluation of parametric data we performed a two-tailed paired *t*-test to account for intra-individual variability where each biological sample served as its own control (left eye – control; right eye – treated). For multivariate analysis across >2 treatments, we performed an ANOVA with the associated Tukey Test to evaluate significant differences between multiple groups.

## Results

### Inhibition of SAHH in damaged retinas

We began by applying 3-Deazaneplanocin A hydrochloride (hereafter referred to as DZN) to NMDA-damaged retinas. Numerous proliferating MGPCs are known to form in NMDA-damaged chick retinas (Fischer and Reh, 2001; Fischer et al., 2004). DZN is a 3-deazaadenosine homolog that inhibits S-adenosylhomocysteine hydrolase (SAHH) which, in turn, increases intracellular levels of SAH and, thereby, blocks that activity of Histone Methyltransferases (HMTs) (Miranda et al., 2009; Tan et al., 2007). DZN-treatment is thought to remove repressive histone methylation and, thus, permit reactivation of genes that are not silenced by DNA methylation (Miranda et al., 2009). Further, DZN has been shown to effectively block the proliferation of cancerous cells (reviewed by (Chase and Cross, 2011; Duan et al., 2020; Herviou et al., 2016). Cellular proliferation is an integral step of MG becoming progenitor-like cells (Fischer and Bongini, 2010; Fischer and Reh, 2003). We found that DZN potently suppressed numbers of proliferating MGPCs that accumulated EdU or expressed PCNA, pHisH3 or neurofilament compared to numbers of MGPCs in control retinas (Fig. 1a-d). Similarly, DZN-treatment suppressed the reactivity (upregulation of CD45), accumulation (DAPI^+^/CD45^+^), and proliferation (EdU^+^/CD45^+^) of microglia (Fig. 1e-g). DZN had no significant effect upon Non-astrocytic Inner Retinal Glial (NIRG) cells. (Supplemental Fig. 1a,b). NIRG cells are a distinct type of glial cell that has been described in the retinas of birds (Fischer et al., 2010; Rompani and Cepko, 2010) and some types of reptiles (Todd et al., 2016b).

**Figure 1.**
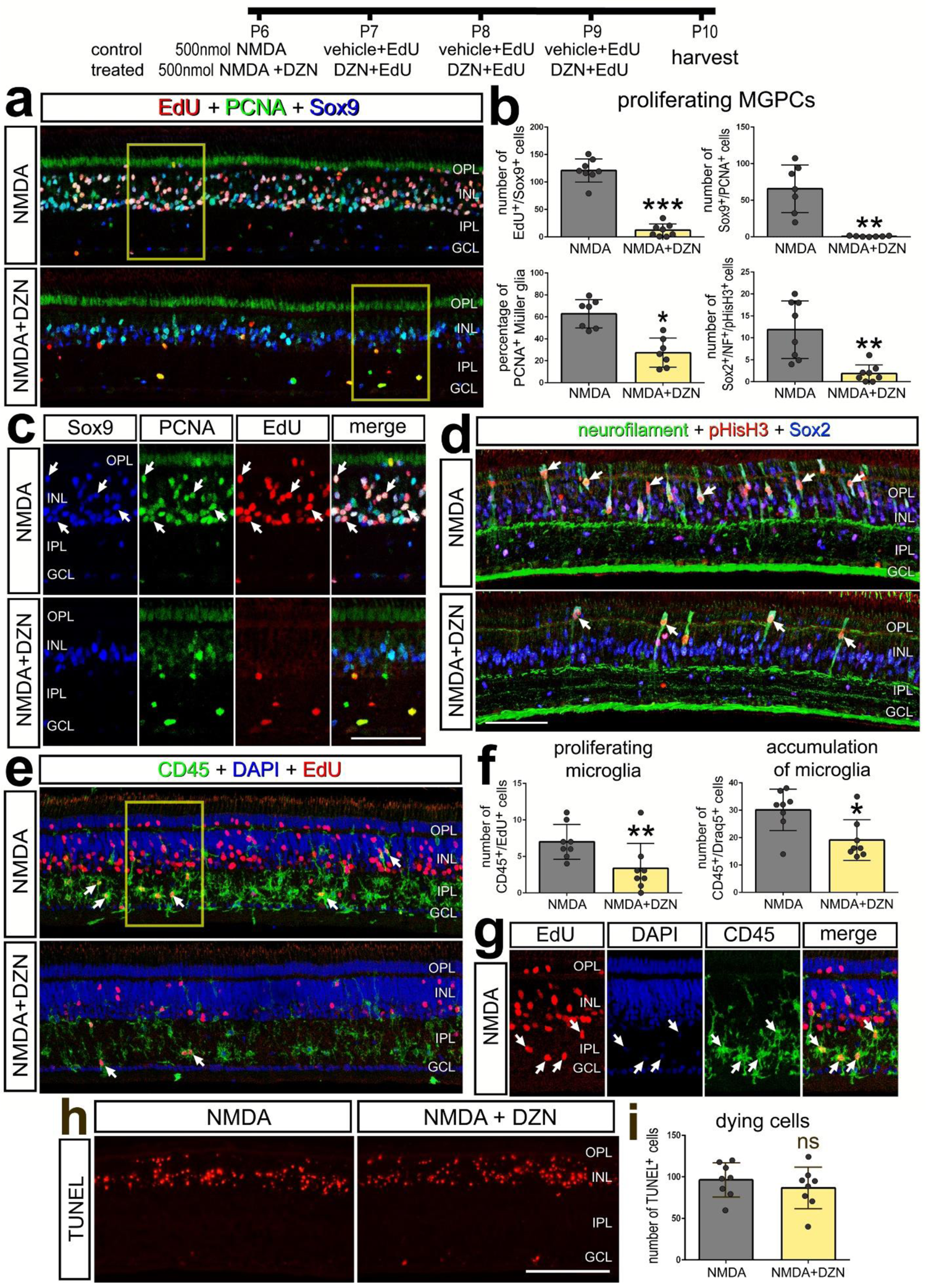
Inhibition of SAAH suppresses the proliferation of MGPCs and microglia. Eyes were injected NMDA ± DZN (SAHH inhibitor) at P6, EdU ± DZN at P7, P8 and P9, and retinas harvested at P10. Sections of the retina were labeled for EdU (red; **a,c,e,g**), PCNA (green; **a,c**), Sox2 (blue; **a,c**), Sox9 (green; **d**), CD45 (**e,g**), DAPI (blue; **e,g**) and TUNEL (dying cells; **h**). Histograms represent the mean (±SD) and each dot represents on biological replicate for proliferating MGPCs (**b**), proliferating and accumulating microglia (**f**), and dying cells (**i**). Significance of difference (*p<0.05, **p<0.001, ***p<0.0001) was determined using a paired t-test. Arrows indicate the nuclei of double- or triple-labeled cells. Abbreviations: ONL – outer nuclear layer, INL – inner nuclear layer, IPL – inner plexiform layer, GCL – ganglion cell layer, ns – not significant.

The effects of DZN were not secondary to influencing cell death. DZN had no effect upon numbers of TUNEL^+^ dying cells in NMDA-damaged retinas (Fig. 1h,i). Further, we found that 3 consecutive daily intraocular injections of DZN alone had no effects upon the reactivity of MG (not shown) or cell death (Supplemental Fig. 1c). Total numbers of Sox2^+^ MG nuclei were significantly reduced in NMDA-damaged retinas treated with DZN, but this was not significantly different from numbers seen in undamaged, saline-treated retinas (Supplemental Fig. 1d,e), consistent with the notion that DZN blocks the proliferation of MGPCs without killing MG. To test whether DZN influenced neuronal differentiation, we applied DZN starting a 3 days after NMDA following the proliferation of MGPCs to avoid the proliferation suppressing effects of DZN. DZN-treatment had no significant effect on the number of neurons produced by MGPCs (data not shown). Collectively, these finding suggest that inhibition of SAHH in damaged retinas potently suppresses the formation of proliferating MGPCs without destroying Müller glia or influencing neuronal cell death.

### Histone methylation in retinas treated with DZN

We next investigated whether DZN-treatment influenced histone methylation by applying antibodies to Histone H3 K27 trimethyl (H3K27me3). H3K27me3 is associated with transcriptional silencing (Cheung and Lau, 2005; Margueron et al., 2005). Immunofluorescence for H3K27me3 was observed in most, if not all, retinal cells. Compared to levels of H3K27me3 immunofluorescence in the nuclei of resting MG in undamaged retinas, levels were not significantly different in the nuclei of activated MG in damaged retinas at 48hrs after treatment (not shown). Levels of H3K27me3 were significantly reduced in area, average intensity per nuclei, and intensity sum in MG nuclei treated with DZN (Fig. 2a-d). Levels of H3K27me3 appeared reduced in outer retinal layers following treatment with DZN (Fig. 2a). Indeed, the average level of H3K27me3 per nuclei and intensity sum for the whole retina was significantly reduced in NMDA-damaged retinas treated with DZN (Fig. 2e,f). The DZN-induced decrease in density sum for H3K27me3 remained significant when subtracting the H3K27me3 immunofluorescence in the nuclei of MG (Fig. 2g).

**Figure 2.**
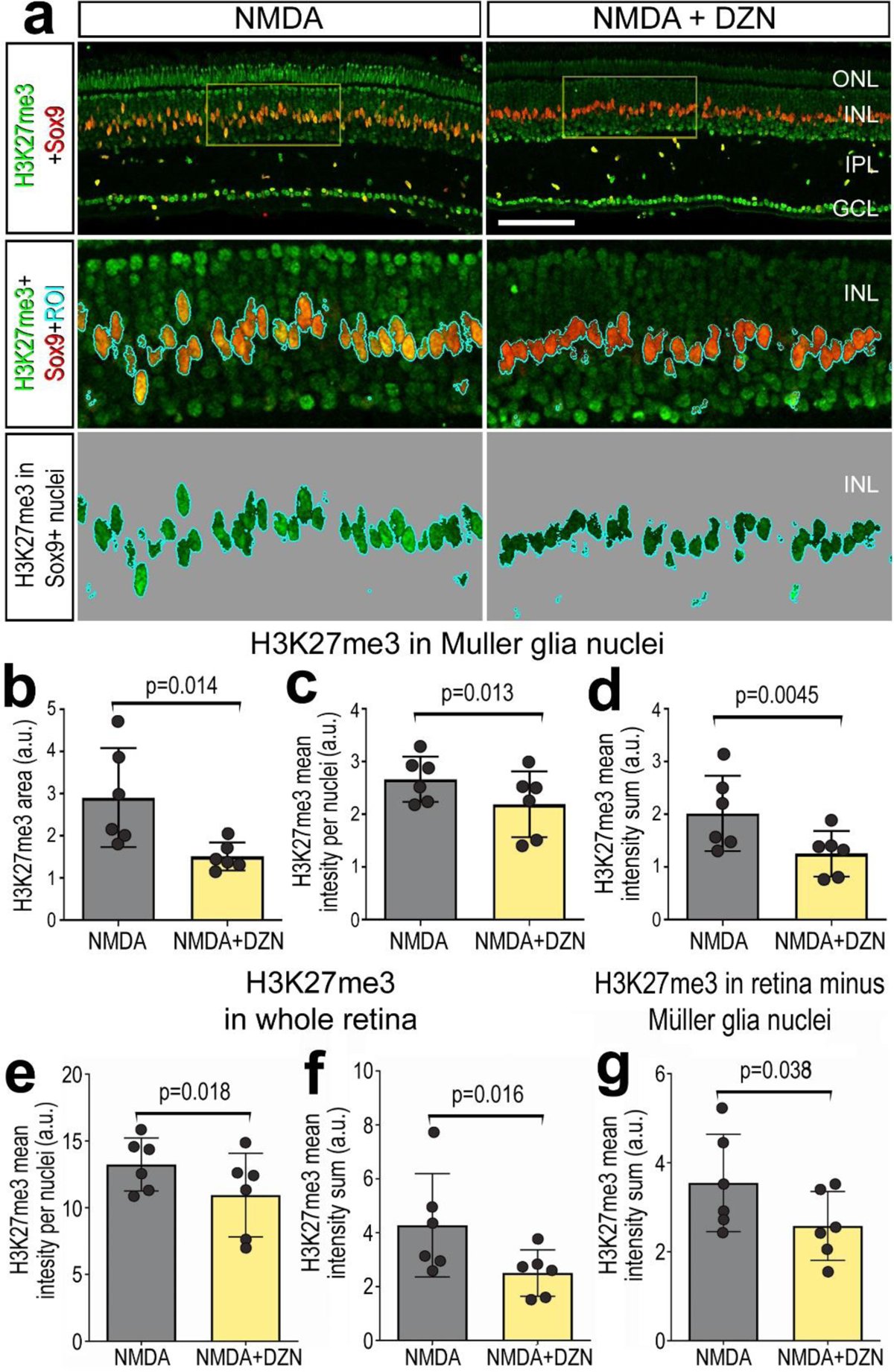
Treatment of retinas with SAHH inhibitor reduces levels of H3K27me3 in the nuclei of MG. Eyes were injected NMDA ± DZN (SAHH inhibitor) at P6, EdU ± DZN at P7 and P8, and retinas harvested at 4hrs after the last injection. Sections of the retina were labeled for Sox9 (red; **a**) and H3K27me3 (green; **a**). The area boxed-out in yellow in the upper panels is enlarged 3-fold in panels below (**a**) The area occupied by Sox9 (MG nuclei) was selected and the H3K27me3 immunofluorescence cut and projected onto a greyscale background for quantification (**a**). Histograms represent the mean (±SD) and each dot represents on biological replicate for H3K27me3 levels in MG nuclei (**b-d**) or total retina (**e-g**). Levels of H3K27me3 were made for for area (**b**), mean intensity per nuclei (**c,e**), density sum in the nuclei of MG (**d, f**) or whole retina minus NG nuclei (**g**). Significance of difference was determined using a paired t-test. Abbreviations: ONL – outer nuclear layer, INL – inner nuclear layer, IPL – inner plexiform layer, GCL – ganglion cell layer. The calibration bar in **a** represent 50 µm.

### Patterns of expression of *AHCY*, *AHCYL1, AHCYL2* and *HMTs*

We analyzed scRNA-seq databases of control and NMDA-damaged chick retinas, for patterns of expression of *AHCY*, *AHCYL1, AHCYL2* and HMTs. We probed libraries of chick retinas from undamaged retinas treated with saline or NMDA at 3, 12 and 48 hrs after treatment, as described previously (Campbell et al., 2021c; Campbell et al., 2022a). After excluding cells with low numbers of genes/cells, putative doublets (high numbers of genes/cells) and dying cells (high percentage of mitochondrial genes), we analyzed a total of 42,202 cells including a total of 4,700 MG (Fig. 3a,b). UMAP ordering of cells resulted in discrete clusters of different types of neurons according and MG according to time after NMDA-treatment (Fig.3a,b). MG were identified based of expression of *VIM*, resting MG (*GLUL*) and activated MG were identified based on expression of *MDK* and *TGFB2* (Fig. 3c,d). We probed for levels and patterns of expression in UMAP heatmap, violin and dot plots (Fig. 3e-h). *AHCY* and *AHCYL1* were expressed by most types of retinal neurons and glia, and levels were significantly different when measured across the whole retina (Fig. 3e,f,h). By comparison, *AHCYL2* was predominantly expression by MG, bipolar cells and some amacrine cells (Fig. 3g). *AHCY* was not highly expressed by MG, whereas *AHCYL1* was widely expressed in MG but levels were not significantly changed after NMDA-treatment (Fig. 3e,f,h). By comparison, *AHCYL2* was widely expressed by MG, and levels and percent expression were increased at 3 hours, decreased at 12 hours and further decreased at 48 hours after NMDA-treatment (Fig. 3g,h). Although many bipolar and amacrine cells expressed *AHCY*, *AHCYL1* and *AHCYL2* (Fig. 3e-g), levels of these genes were not significantly changed after NMDA-treatment, with the exception of *AHCLY1* at 3 hrs after treatment (Fig. 3h).

**Figure 3.**
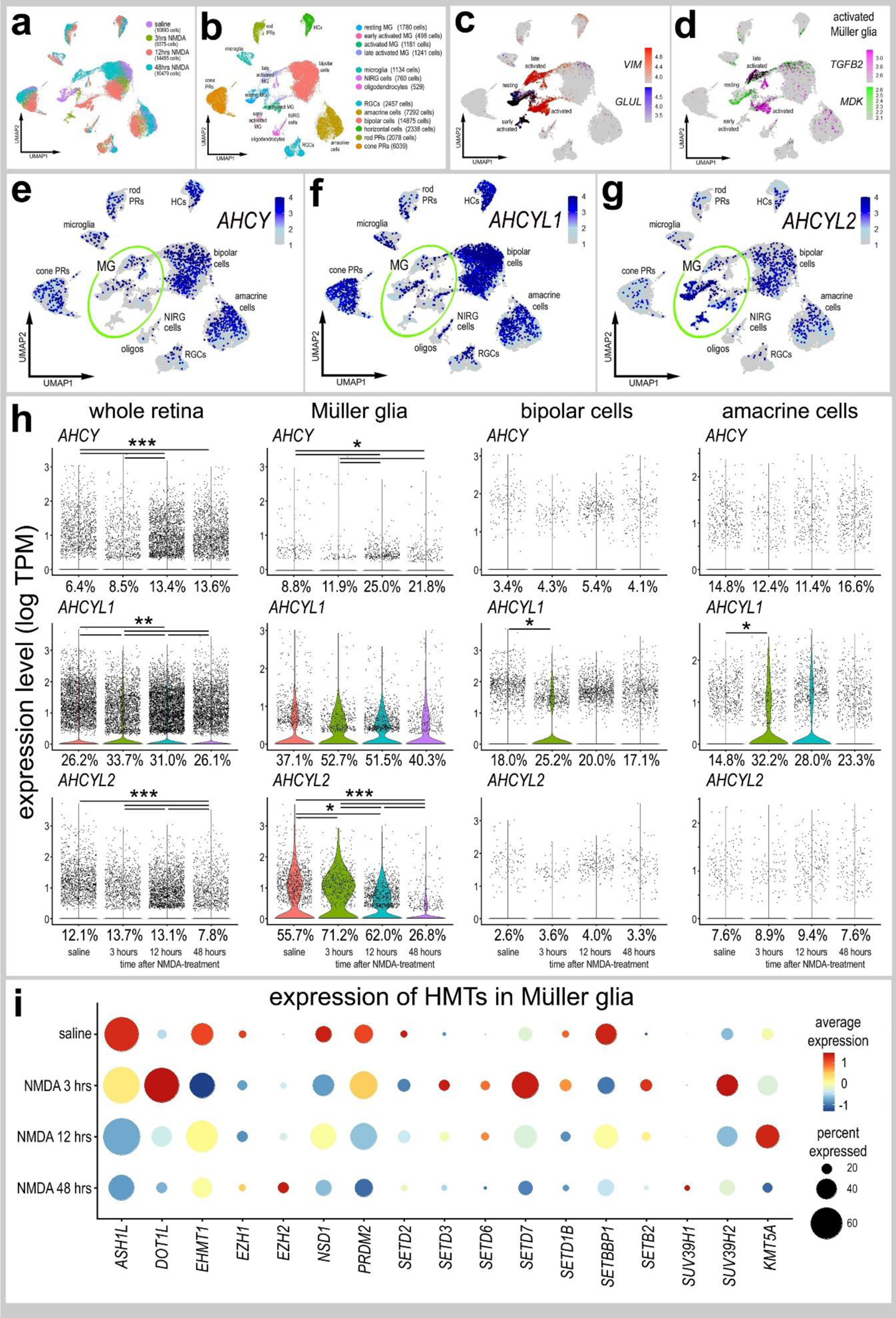
Patterns of expression of *ACHY, ACHYL1* and *ACHYL2* in normal and damaged retinas. scRNA-seq was used to identify patterns and levels of expression of *ACHY, ACHYL1, AHCYL2* and HMTs in control and NMDA-damaged retinas at 3, 12 and 48 hours after treatment (**a**). UMAP clusters of cells were identified based on well-established patterns of gene expression (see Methods; **b**). MG were identified by expression of *VIM* and *GLUL* in resting MG (**c**), and *TGFB2* and *MDK* activated MG (**d**). Each dot represents one cell and black dots indicate cells that express 2 or more genes (**c**-**d**). The violin plots in **h** illustrate the expression levels and percent expressing cells for *ACHY, ACHYL1* and *AHCYL2* whole retina, Müller glia, bipolar cells and amacrine cells in control and NMDA-damaged retinas at 3, 12 and 48 hours after treatment. The dot plot in **i** illustrates average expression (heatmap) and percent expressed (dot size) for different HMTs in MG that are significantly (p<0.01) up- or downregulated at different times following NMDA-treatment. Significant of difference (*p<0.01, **p<10^-10^, ***p<10^-20^) was determined by using a Wilcox rank sum test with Bonferroni correction.

Expression levels of HMTs in MG are illustrated in a dot plot (Fig. 3i). We found that many HMTs, including *ASH1L, EMT1, EZH1, NSD1, SETD2, SETD1B, SETBP1* and *SUV39H2* were significantly decreased in MG following NMDA-treatment (Fig. 3i). By comparison, some HMTs, including *DOT1L, SETD3, SETD6, SETD7, SETB2* and *KMT5A*, were rapidly upregulated by MG between 3 and 12 hours after NMDA-treatment (Fig. 3i). Among HMTs, only EZH2 and SUV39H1 were upregulated by MG at 48 hours after NMDA (Fig. 3i), when proliferating MGPCs are known to form (Fischer and Reh, 2001).

### Chromatin access and gene expression following inhibition of AHCYs

We treated normal and NMDA-damaged retinas treated with vehicle or DZN and harvested retinal cells to generate scRNA-seq and scATAC-seq libraries. We aggregated scRNA-seq libraries from retinas treated with saline, saline+DZN, NMDA and NMDA+DZN (Fig. 4a,b). DZN-treatment resulted in distinct clustering of MG from the different treatment groups, whereas the UMAP-ordered retinal neurons from different treatments clustered together (Fig. 4b). Resting MG, activated MG and MGPCs were identified based on distinct patterns of expression of *RLBP1, GLUL, SFRP1* (resting), *TGFB2, MDK, PMP2* (activated), and *TOP2A, PCNA, SPC25, CDK1, CCNB2* (MGPCs) (Fig. 4c-e, g-j). The cluster of resting MG was comprised primarily of MG from saline-treated retinas, whereas DZN-treated MG formed a distinct cluster (Fig. 4g-i). The cluster of MGPCs was comprised almost exclusively of MG from NMDA-treated retinas, whereas activated MG was comprised of cells from retinas treated with DZN-saline, DZN-NMDA and NMDA alone (Fig. 4g-i). The cluster of MGPCs correlated well with Cell-Cycle Scoring for cells in S-phase and G2/M-phase (Fig. 4k). We identified numerous differentially expressed genes (DEGs) that were significantly up- or down-regulated in MG in different treatment groups (Fig. 4f). DZN-treatment of MG in undamaged retinas significantly upregulated genes associated with resting MG, glial transcription factors, and cell signaling pathways known to influence MGPC formation (Fig. 4l), including MAPK-(*FGF9, EGF*) (Fischer et al., 2009a; Fischer et al., 2009b), BMP-(*BMPR2*) (Todd et al., 2017), and Wnt-signaling (*WIF1, SFRP1, SFRP2*) (Gallina et al., 2016). By comparison, DZN-treatment of MG in undamaged retinas downregulated genes associated with cell signaling pathways known to influence MGPC formation (Fig. 4l), including midkine/pleotrophin (*MDK, PTN, SDC4*) (Campbell et al., 2021b), Jak/Stat-signaling (*SOCS3*) (Todd et al., 2016a), Notch-signaling (*MAML2*) (Ghai et al., 2010; Hayes et al., 2007) and MMPs (*TIMP2*) (Campbell et al., 2019). DZN-treatment of MG in NMDA-damaged retinas significantly upregulated transcription factors that convey glial identity, such as *VSX2* and *HES1* (Furukawa et al., 2000; Wall et al., 2009), and genes associated with cell signaling pathways, such as MAPK (*MAPK6, SPRY2*) and Wnt/β-catenin (*CTNNB1*) (Fig. 4m). DZN-treatment of MG in NMDA-damaged retinas significantly downregulated genes associated with chromatin modifications (including *EZH2* and several HMG’s), cell cycle progression, transcription factors, and cell signaling factors, particularly those related to TGFβ/BMP-signaling (which is known to regulate the formation of MGPCs in the chick retina) (Fischer et al., 2004; Todd et al., 2017) (Fig. 4m).

**Figure 4.**
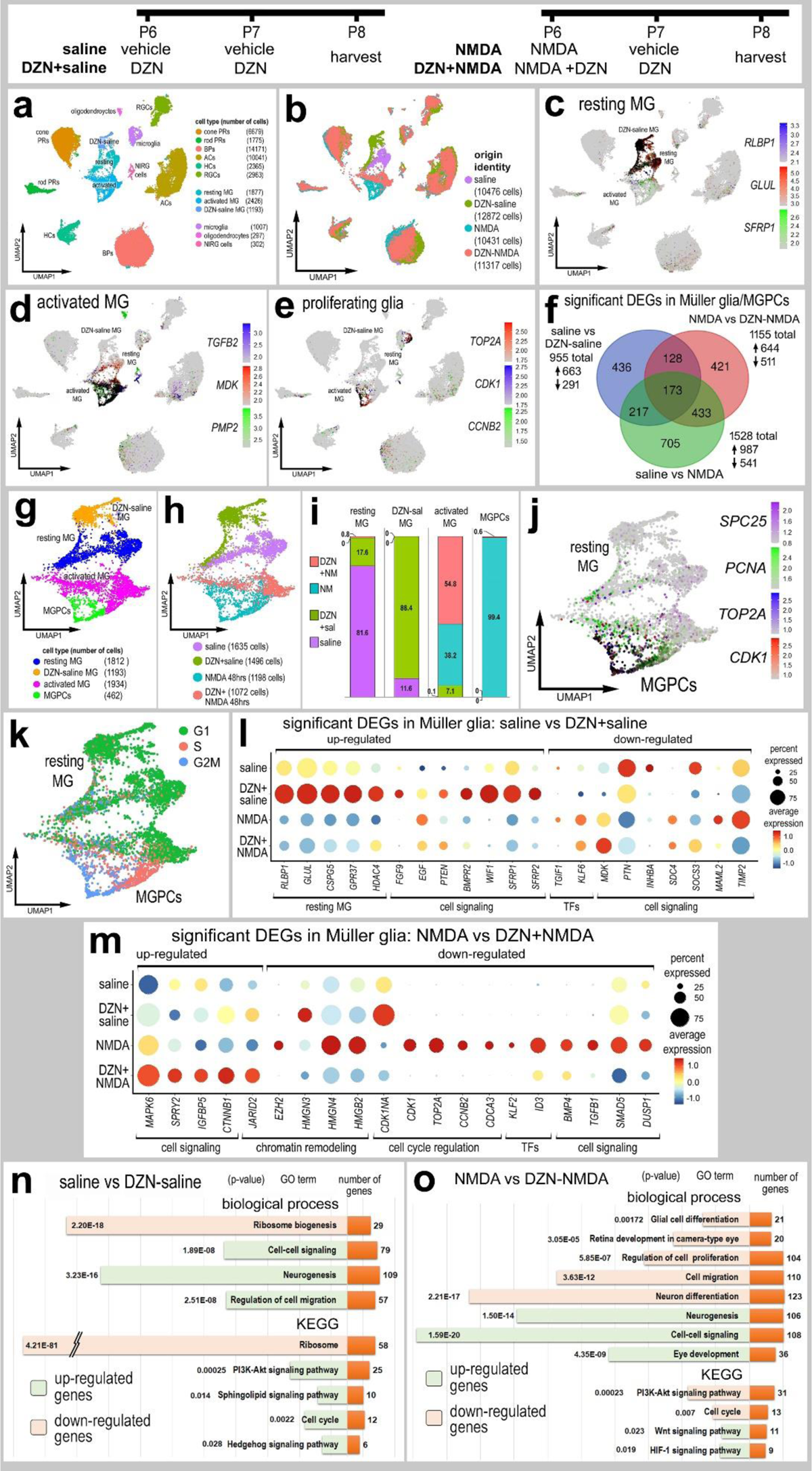
Treatment of retinas with SAHH inhibitor and NMDA influences gene expression in MG. Retinas were treated with saline ± DZN or NMDA ± DZN, and scRNA seq libraries were generated. UMAP plots were generated and clusters of cells identified based on established markers (**a**,**b**). MG were identified based on expression of genes associated with resting glia (**c**), activated glia (**d**) and proliferating MGPCs (**e**). Lists of DEGs were generated (supplemental table 1) for MG from retinas treated with saline vs DZN-saline, saline vs NMDA, and NMDA vs DZN-NMDA and plotted in a Venn diagram (**f**). UMAP clustered MG were isolated and analyzed for occupancy according to library of origin (**g**,**h**,**i**). MGPCs were identified based on high levels of expression of *SPC25, PCNA, TOP2A* and *CDK1* (**j**). Dot plots illustrate the percentage of expressing MG (size) and significant (p<0.001) changes in expression levels (heatmap) for genes in MG from retinas treated with saline vs saline-DZN (**k**) and NMDA vs NMDA-DZN (**l**). Genes related to resting MG, cell signaling, chromatin remodeling, cell cycle regulation or transcription factors (TFs) were up- or down-regulated (**k**,**l**). GO enrichment analysis was performed for lists of DEGs in MG for retinas treated with saline ± DZN and NMDA ± DZN (**n,o**). Gene modules for upregulated (green) and downregulated (peach) genes were grouped by GO category with P values and numbers of genes for each category (**n,o**).

We performed gene ontology (GO) enrichment analyses for DEGs in MG resulting from SAHH inhibition in normal and damaged retinas. In DZN-treated MG in undamaged retinas, we observed upregulated DEGs in gene modules associated with neurogenesis, cell migration, cell cycle and cell signaling, namely PI3K-Akt, sphingolipids and the Hedgehog pathway (Fig. 4n). Downregulated DEGs occupied gene modules associated with ribosome biogenesis (Fig. 4n). In DZN-treated MG in damaged retinas, we observed upregulated DEGs in gene modules associated with neurogenesis, eye development, and cell signaling including Wnt and HIF-1 pathways (Fig. 4o). Downregulated DEGs occupied gene modules associated with glial/neuronal differentiation, cellular proliferation, retinal development, cell migration and PI3K-Akt signaling (Fig. 4o).

Following scATAC-seq UMAP embedding of cells revealed distinct clusters of different types of neurons and glia (Fig. 5a,b). We identified clusters of cell types from elevated chromatin access for genes known to be expressed by MG (*SOX2, NOTCH1, PAX6*), oligodendrocytes and NIRG cells (*NKX2-2, OLIG2*), amacrine cells (*TFAP2A, PAX6*), horizontal cells (*PAX6*), bipolar cells (*OTX2*, *VSX2*), ganglion cells (*ISL2*), cone photoreceptors (*ISL2*, *CALB1*, *GNAT2*), and rod photoreceptors (*RHO*) (Fig. 5c). DZN- and NMDA-treatments had little effect upon UMAP-clustering of retinal neurons, whereas MG from different treatments were spatially separated (Fig. 5a,b,d), indicating that changes in chromatin access resulting from NMDA-treated or inhibition of SAHH are primarily manifested in MG, but not other types of retinal cells. Unfortunately, very few MG (94 cells) from NMDA+DZN-treated retinas were captured (Fig. 5a), making statistical analysis impossible.

**Figure 5.**
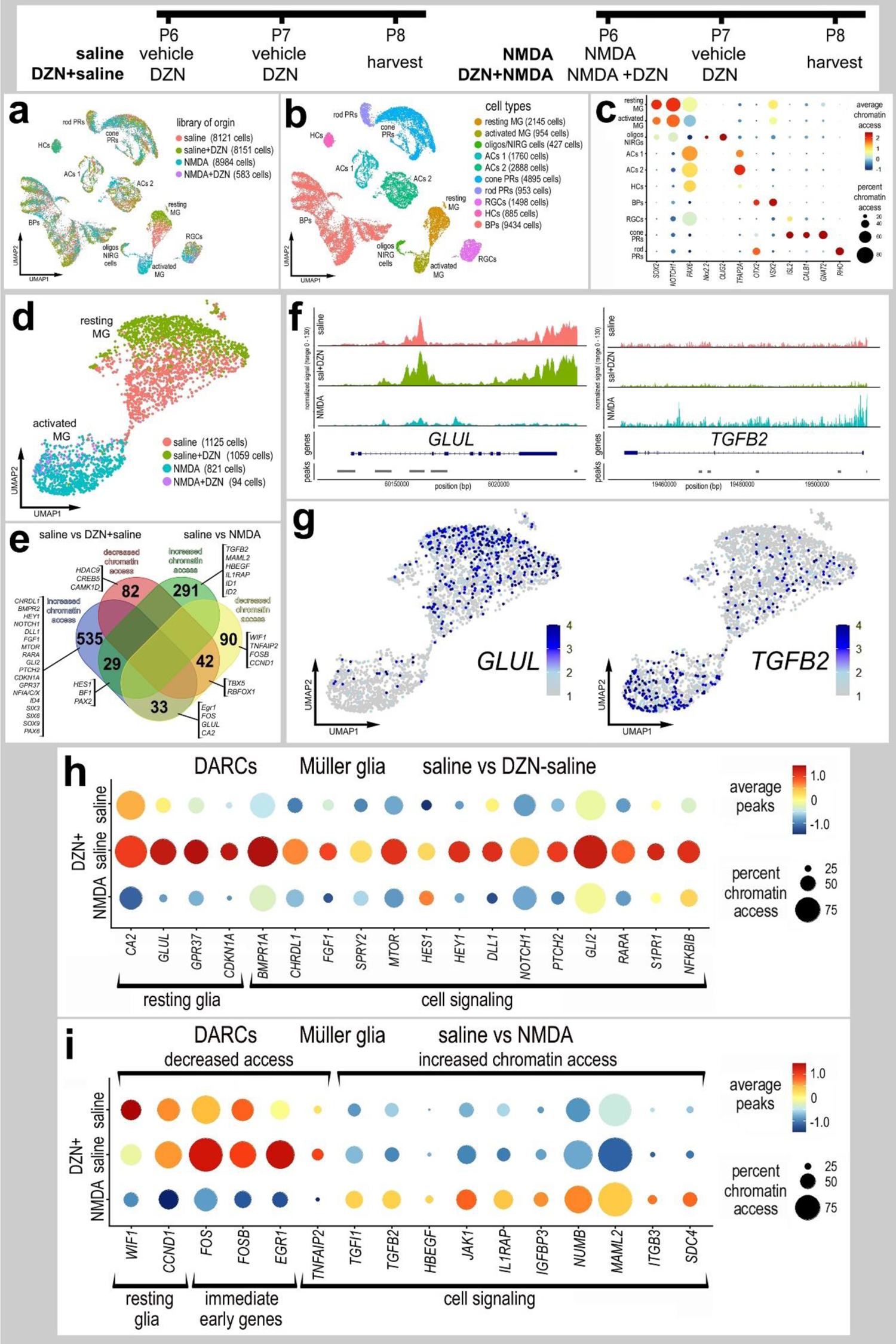
Treatment of retinas with SAHH inhibitor influences chromatin access in MG. Eyes were injected with saline ± DZN or NMDA ± DZN, retinas were harvested 24hrs after the last injection and scATAC-seq libraries were generated (**a**). UMAP plots were generated and clusters of cells identified based on chromatin access for genes known to be associated with retinal cell types (**b**,**c**). Genes with enhanced chromatin access in distinct cells types included *GNAT2* (cone PRs), *ISL2* and *POU4F2* (RGCs), *VSX2* (bipolar cells), *PAX6* (amacrine and horizontal cells), *TFAP2A* (amacrine cells), and illustrated in a dot plot (**c**). MG were isolated and re-ordered in UMAP plots (**d**). UMAP clusters of MG formed for resting MG (saline and saline+DZN-treated cells) and activated MG (NMDA and NMDA+DZN-treated cells) (**d**). Lists of genes with differential accessible chromatin regions (DACRs; supplemental tables 2 and 3) were generated and a The Venn diagram (**e**) constructed to illustrate numbers of genes with significantly up- and down-regulated chromatin access according to treatment. Significance of difference (p<0.05) was determined by using a Wilcox rank sum test with Bonferroni correction. Clusters or resting and activated MG were distinguished based on differential chromatin access to genes such as *GLUL, TGFB2* (**f,g**)*, EGR1, FOSB, FOS, CDKN1A* and *HEYL* (**h,i**).

There were significant changes in genes with differentially accessible regions of chromatin (DARC) when comparing scATAC-seq for MG from saline±DZN and NMDA-treated retinas (Fig. 5e-k). Treatment with DZN or NMDA resulted primarily in increased chromatin access in numerous genes, whereas decreased chromatin access occurred for relatively few genes (Fig. 5e). For example, DZN-treatment increased, while NMDA decreased, chromatin access for immediate early genes, such as *FOS, FOSB* and *EGR1*, and markers of resting glia, such as *CDK1NA, CA2* and *GLUL* (Fig. 5e-h). DZN-treatment of undamaged retinas resulted in changes in chromatin access in genes associated with maintaining glial phenotype that are expressed by resting MG such as *GPR37* and *CDKN1A* (Fig. 5e,h). In addition, DZN-treatment of undamaged retinas resulted in changes in chromatin access for many genes that have been implicated in regulating the formation of MGPCs, including signaling pathways for BMP and TGFβ (*TGIF1, TGFB2, BMPR1A, CHRDL1*) (Todd et al., 2016a), MAPK-signaling (*FGF1, SPRY2*) (Fischer et al., 2009a; Fischer et al., 2009b), Notch (*HEY1, HES1, DLL1, NOTCH1*) (Ghai et al., 2010; Hayes et al., 2007), Hedgehog (*GLI2, PTCH2*) (Todd and Fischer, 2015), (Gallina et al., 2014b), retinoic acid (*RARA*) (Todd et al., 2018), mTor (*MTOR)* (Zelinka et al., 2016), and pro-inflammatory signaling (*NFKBIB, S1PR1)* (Fig. 5e-h). MG from NMDA-damaged retinas had increased chromatin access to many different genes including different transcription factors and cell signaling pathways, particularly those associated with TGFβ-signaling (Fig. 5e-g, i). In addition, MG from NMDA-damaged retinas had decreased chromatin access for immediate early genes which are known to be upregulated by activated MG (Fischer et al., 2009a; Fischer et al., 2009b), and genes associated with signaling pathways known to influence the formation of MGPCs including HBEGF (Todd et al., 2015), gp130/JAK/STAT (*JAK1, IL1RAP*) (Todd et al., 2016a), Notch (*NUMB, MAML2*) (Ghai et al., 2010; Hayes et al., 2007), and midkine/pleotrophin (*ITGB3, SDC4*) (Campbell et al., 2021b) (Fig. 5e,i).

We next compared the DEGs from scRNA-seq and DARCs from scATAC-seq in MG in undamaged retinas treated with SAHH inhibitor. We found 127 genes with increased expression and increased chromatin access (Fig. 6a), and these genes included pro-glial transcription factors (*HES1, ID4, NFIX*) and genes highly expressed by resting glia (*GLUL, GPR37, CSPG5, CDKN1A*). We found 21 genes with increased mRNA levels and decreased chromatin access (Fig. 6a), and these genes included those upregulated in reactive glia (*GFAP, VIM*) and different cell signaling pathways known to influence MG (*FSTL1, CHRDL1, FOS, SOCS3, TIMP2*) (Campbell et al., 2019; Gallina et al., 2016; Todd et al., 2016a). There were only 2 genes with decreased mRNA and decreased chromatin access (Fig. 6a).

**Figure 6.**
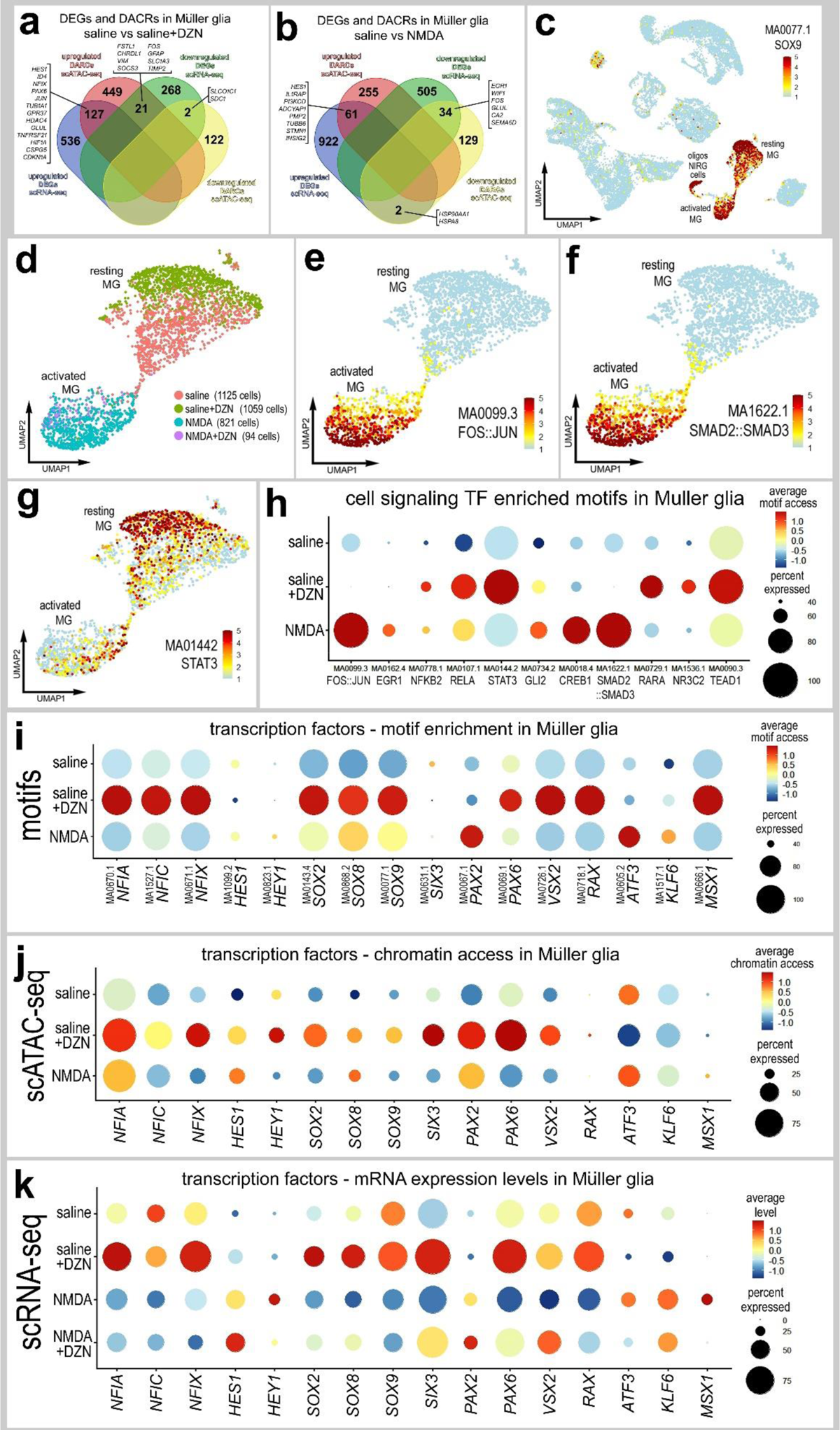
Comparison of differentially expressed genes (DEGs), differentially accessible chromatin regions (DACRs) and access to transcription factor binding motifs in MG. Venn diagrams illustrate numbers of genes with increased or decreased expression levels or chromatin access in MG from retinas treated with saline vs saline+DZN (**a**) or saline vs NMDA (**b**). UMAP heatmap plot in **c** illustrates access to SOX9 DNA-binding motifs across all types of retinal cells. Motifs enriched in activated and resting MG were identified, and included motifs for transcriptional effectors of cell signaling pathways (**d-h**). UMAP heatmap plots for MG illustrate differential access to motifs for FOS::JUN, SMAD2::SMAD2 and STAT3 in resting and activated MG (**e-g**). Dot plots illustrate heatmaps for levels of access and percent of cells with access for transcriptional effectors of cell signaling pathways (**h**) or transcription factors (**i**). For direct comparison with motif access, dot plots illustrate levels of chromatin access (**j**) and mRNA (**k**) for key transcription factors.

We next compared the DEGs from scRNA-seq and DARCs from scATAC-seq in MG in saline vs NMDA-damaged retinas. We found 61 genes with increased expression and increased chromatin access (Fig. 6b), and these genes included a pro-glial transcription factor (*HES1*), and genes associated with cell-signaling (*ILRAP1, PI3KCD, ADCYAP1, INSIG2*) and activated glia (*PMP2, TUBB6*). We found 34 genes with decreased mRNA and decreased chromatin access (Fig. 6b), and these genes included those highly expressed by resting MG (*WIF1, GLUL, CA2*) and immediate early genes (*FOS, EGR1*). There were only 2 genes with increased mRNA and decreased chromatin access (Fig. 6b). We were not able to identify genes from scRNA-seq and scATAC-seq in NMDA-damaged retinas treated with SAHH-inhibitor because too few MG were captured in the scATAC libraries.

We next identified transcription factor DNA-binding motifs with differential chromatin access in promoter regions in MG. MG were identified based on significant access to motifs such as *SOX9* (Fig. 6c), *RAX, MSX1* and *VSX2* (not shown). Activated and resting MG were distinguished based of enriched access for motifs for transcriptional effectors of different cell signaling pathways such as *FOS::JUN*, *SMAD2::SMAD3* and *STAT3* (Fig. 6e-g). In addition, we identified significant differences in enriched access to motifs related to NFkB (*NFKB2, RELA*), Hedgehog (*GLI2*), retinoic acid (*RARA*), glucocorticoid (*NR3C2*) and Hippo-signaling pathways (*TEAD1*), which are known to influence the formation of MGPCs (Gallina et al., 2014b; Palazzo et al., 2020; Palazzo et al., 2022; Rueda et al., 2019; Todd and Fischer, 2015; Todd et al., 2018; Young et al., 2000). We next analyzed levels of mRNA (scRNA-seq), chromatin access (scATAC-seq) and enriched motif access in MG for transcription factors that are known to be important for retinal development and the formation of MGPCs. We observed consistent increases across levels of mRNA, chromatin access and motif enrichment in DZN-treated MG in undamaged retinas for *NFIA/C/X*, *SOX2/8/9*, *PAX6, VSX2* and *RAX* (Fig. 6i-k). By comparison, we observed increases across levels of mRNA, chromatin access and motif enrichment in NMDA-treated MG for *HES1, HEY1, PAX2, ATF3* and *KLF6* (Fig. 6i-k).

### Ligand-Receptor interactions among MG and microglia

We next bioinformatically isolated MG and microglia, re-embedded these cells in UMAP plots and probed for cell signaling networks and putative ligand-receptor interactions using SingleCellSignalR (Cabello-Aguilar et al., 2020). We focused our analyses on MG and microglia because there is significant evidence to indicate autocrine and paracrine signaling among these cells in the context of glial reactivity and the formation of MGPCs (Fischer et al., 2014; White et al., 2017; Zelinka et al., 2012). UMAP plots revealed distinct clusters of resting MG (*GLUL, GPR37, RLBP1*) from saline and saline-DZN libraries, activated MG (*MDK, HBEGF, TGFB2*), MGPCs (*TOP2A. SPC25, CDK1*) and a cluster of microglia (*C1QA, SPI1, CSF1R*) (Figs. 7a-f). Numbers of LR-interactions (significant expression of ligand and receptor) between cell types in the different treatment groups varied from 124 to 292 (Fig. 7g). We identified the 40 most significant paracrine LR-interactions for MG to microglia and microglia to MG for each treatment group (Figs. 7h-k). We next identified paracrine LR-interactions between glia unique to saline vs saline+DZN treatment groups (Figs. 7l,n). In the saline-treatment group, we found 41 unique paracrine LR-interactions including midkine-, FGF-, TGFβ- and activin-signaling. By comparison, saline+DZN-treatment resulted in 59 unique paracrine LR-interactions between glia such as EGF-, HBEGF-, IGF- and Notch-signaling. In the NMDA-treatment groups (Fig. 7m), we found 30 unique paracrine LR-interactions including FGF- and laminin/integrin-signaling. By comparison, NMDA+DZN-treatment resulted in 29 unique paracrine LR-interactions between glia such as EGF-, HBEGF-, BDNF-vitronectin/integrin- and plasminogen activator urokinase (*PLAU*) - signaling. Many of the above-mentioned pathways have been implicated in the formation of MGPCs in the chick retina including HB-EGF, IGF/FGF/MAPK-, Wnt/-β catenin-, Notch-, TGFβ/Smad- and midkine-signaling (Campbell et al., 2021b; Fischer et al., 2009a; Gallina et al., 2016; Ghai et al., 2010; Hayes et al., 2007; Todd et al., 2015; Todd et al., 2017; Zelinka et al., 2016).

**Figure 7.**
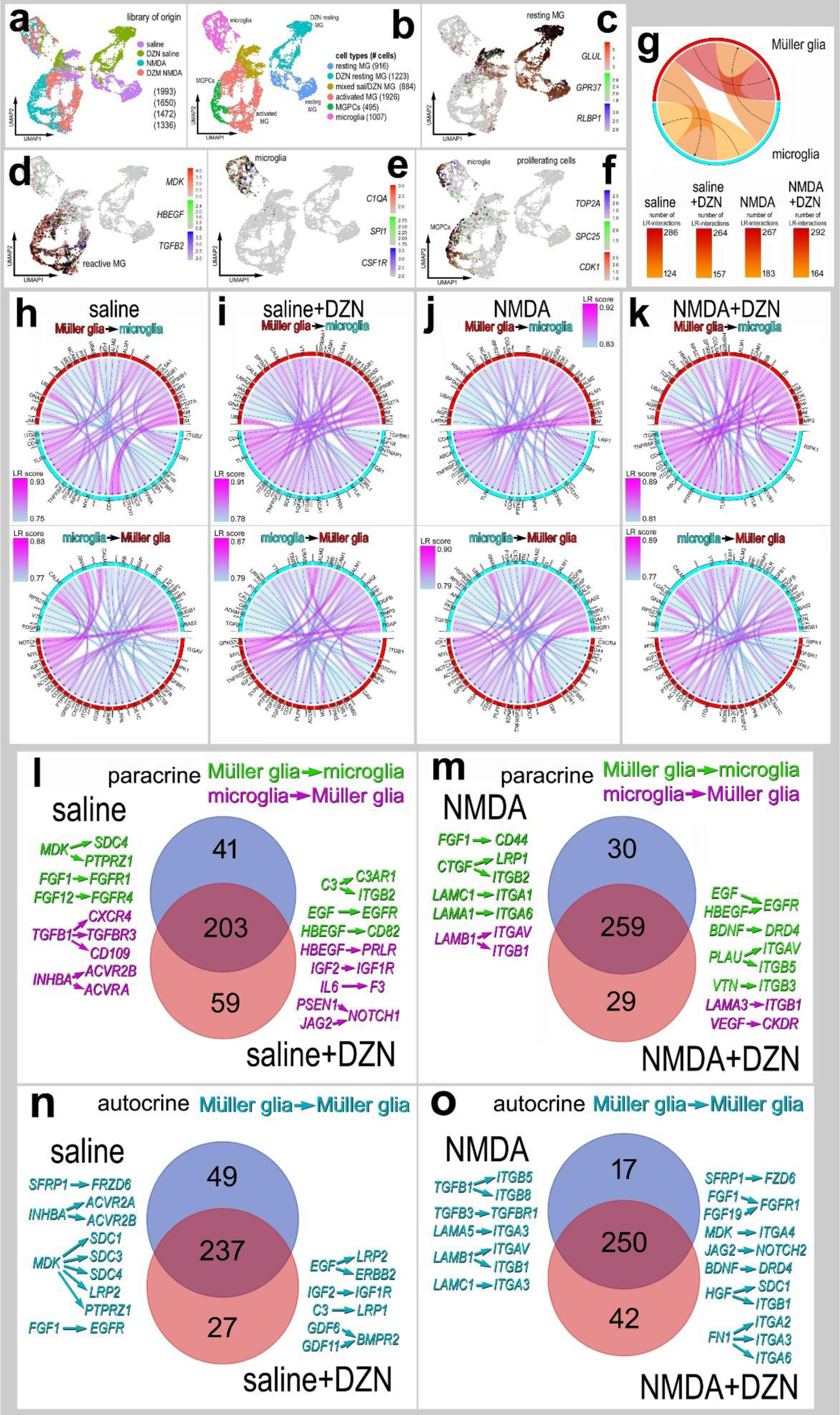
Ligand-receptor (LR) interactions inferred from scRNA-seq data between microglia and MG. Retinal microglia and MG were isolated, re-embedded and ordered in UMAP plots (**a,b**). Treatment groups included saline, saline + DZN, NMDA and NMDA + DZN. Cells were identified based on cell-distinguishing markers: Resting MG - *GLUL, GPR37, RLBP1* (**c**); reactive MG – *MDK, HBEGF, TGFB2* (**d**); microglia - *C1QA, SPI1, CSF1R* (**e**); proliferating cells (microglia and MPGCs) - *TOP2A, SPC25, CDK1* (**f**). Glia from different treatment groups were analyzed using SingleCellSignalR to generate chord diagrams and illustrate numbers of autocrine and paracrine LR-interactions (**g**). The LRscore, with the most significant LR-interactions approaching a value of 1, utilizes the mean normalized read count matrix to regularize the Ligand and Receptor read counts independent of dataset depth. Paracrine LR-interactions were identified for glial cells for different treatment groups including saline (**h**), DZN-saline (**i**), NMDA (**j**) and DZN-NMDA (**k**). For each treatment group, the 40 most significant LR-interactions between microglia and MG were identified and illustrated in chord plots with LR score heat maps (**h-k**). Treatment-specific differences in glial paracrine LR-interactions in saline vs saline + DZN (**l**) and NMDA vs NMDA+DZN (**m**) are illustrated in Venn diagrams with select interactions. Treatment-specific differences in glial autocrine LR-interactions in MG in saline vs saline + DZN (**n**) and NMDA vs NMDA +DZN (**o**) are illustrated in Venn diagrams with select interactions.

We next identified MG-specific autocrine LR-interactions in saline ± DZN and NMDA ± DZN treatment groups (Figs. 7n,o). Unique to the saline-treatment group for autocrine MG, we found 49 unique LR-interactions including midkine-, Wnt-, FGF- and activin-signaling. By comparison, saline+DZN-treatment resulted in 27 unique autocrine MG LR-interactions such as EGF-, IGF- and GDF/BMPR2-signaling. In the NMDA-treatment groups (Fig. 7o), we found 17 unique autocrine MG LR-interactions including TGFβ- and laminin/integrin-signaling. By comparison, NMDA+DZN-treatment resulted in 42 unique autocrine MG LR-interactions between glia such as FGF-, Notch-, Midkine-, Wnt-, BDNF-, Hepatocyte Growth Factor (HGF) /integrin- and fibronectin (*FN1*)/integrin-signaling. Many of the above-mentioned pathways have been implicated in the formation of MGPCs in the chick retina including HB-EGF, IGF/FGF/MAPK-, Wnt/β - catenin-, Notch-, BMP/Smad- and midkine-signaling (Campbell et al., 2021b; Fischer et al., 2009a; Gallina et al., 2016; Ghai et al., 2010; Hayes et al., 2007; Todd et al., 2015; Todd et al., 2017; Zelinka et al., 2016).

### Inhibition of SAHH suppresses mTor signaling in MG in damaged retinas

We next investigated whether the inhibition of SAHH influenced cell signaling pathways that are known to promote the formation of MGPCs. cFos, pS6 (mTor-signaling), pStat3 (Jak/Stat-signaling) and pSmad1/5/8 (BMP/Smad-signaling) are known to be rapidly upregulated in MG after NMDA-treatment and promote the formation of proliferating MGPCs (Fischer et al., 2009a; Todd et al., 2016a; Todd et al., 2017; Zelinka et al., 2016). Thus, we applied NMDA ± DZN at P6, vehicle or DZN at P7, and harvested retinas 4hrs after the last injection. At this early time point there was no significant difference in numbers of dying cells with DZN-treatment (Fig. 8a,b), similar to findings at 72hrs after NMDA (Fig. 1h,i). DZN-treatment had no effect upon damaged-induced increases in cFos, pStat3 or pSmad1/5/8 in MG in damaged retinas (Fig. 8a,c,e,f). By comparison, we found that DZN-treatment resulted in a significant decrease in levels of pS6 in MG in damaged retinas (Fig. 8a,d).

**Figure 8.**
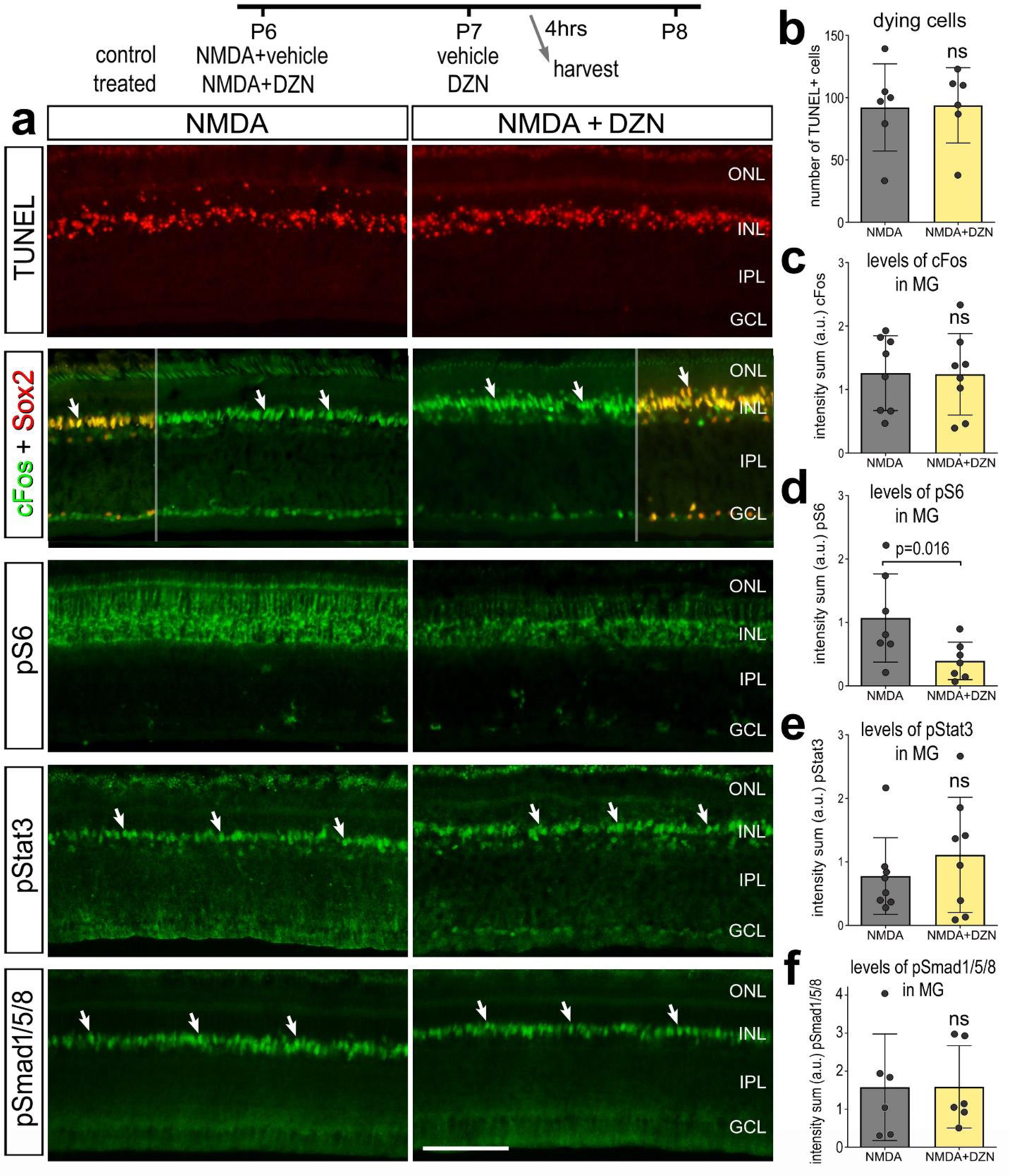
Cell signaling in MG in damaged retinas is influenced by SAHH inhibitor. Eyes were injected with NMDA ± DZN (SAHH inhibitor) at P6, vehicle or DZN at P7, and eyes harvested at 4 hrs after the last injection. Retinal sections were for cell death (TUNEL) or antibodies to cFos and Sox2, pS6 and pStat3 (**a**). The calibration bars in **a** represent 50 µm. (**b**-**f**) Histograms represent the mean (±SD) and each dot represents one biological replicate. Significance of difference (p-value) was determined using a paired t-test. Abbreviations: ONL – outer nuclear layer, INL – inner nuclear layer, IPL – inner plexiform layer, GCL – ganglion cell layer, ns – not significant.

### Inhibition of SAHH in undamaged retinas and CMZ proliferation

To test whether SAHH activity and histone methylation was involved in the formation of MGPCs in the absence of retinal damage we applied a growth factor-treatment paradigm. Three consecutive daily intraocular injections of insulin and FGF2 are known to induce the formation of proliferating MGPCs in the absence of damage (Fischer et al., 2002b; Fischer et al., 2014). There was a significant decrease in the number of EdU/Sox2-positive nuclei (proliferating MGPCs) in insulin+FGF2-treated retinas treated with DZN (Fig. 9b,c). Consistent with these findings, there was a significant decrease in the number of pHisH3/Sox2/neurofilament-positive cells in insulin+FGF2-treated retinas treated with DZN (Fig. 9d,e). In central regions of the retina, insulin+FGF2 stimulated the migration of MG nuclei away from the middle of the INL without EdU incorporation (Fig. 9f,g). The migration of nuclei is correlated with the proliferation of MGPCs (Fischer and Reh, 2001), but migration can occur without re-entry into the cell cycle (Fischer et al., 2014; Todd and Fischer, 2015). Treatment with insulin+FGF2+DZN inhibited the migration of MG nuclei away from the middle of the INL (Fig. 9f,g). The proliferation and reactivity of microglia was unaffected by DZN in insulin+FGF2-treated retinas (not shown). Treatment with insulin and FGF2 is known to stimulate MG to rapidly and transiently upregulate cFos, pSmad1/5/8, pS6 and pStat3 (Fischer et al., 2009c; Todd et al., 2016a; Todd et al., 2017; Zelinka et al., 2016). DZN-treatment had no effects upon patterns of immunofluorescence for cFos, pStat3 or pS6 (Fig. 9h and not shown), whereas immunofluorescence of pSmad1/5/8 was sustained and elevated in the nuclei of MG (Fig. 9h).

**Figure 9.**
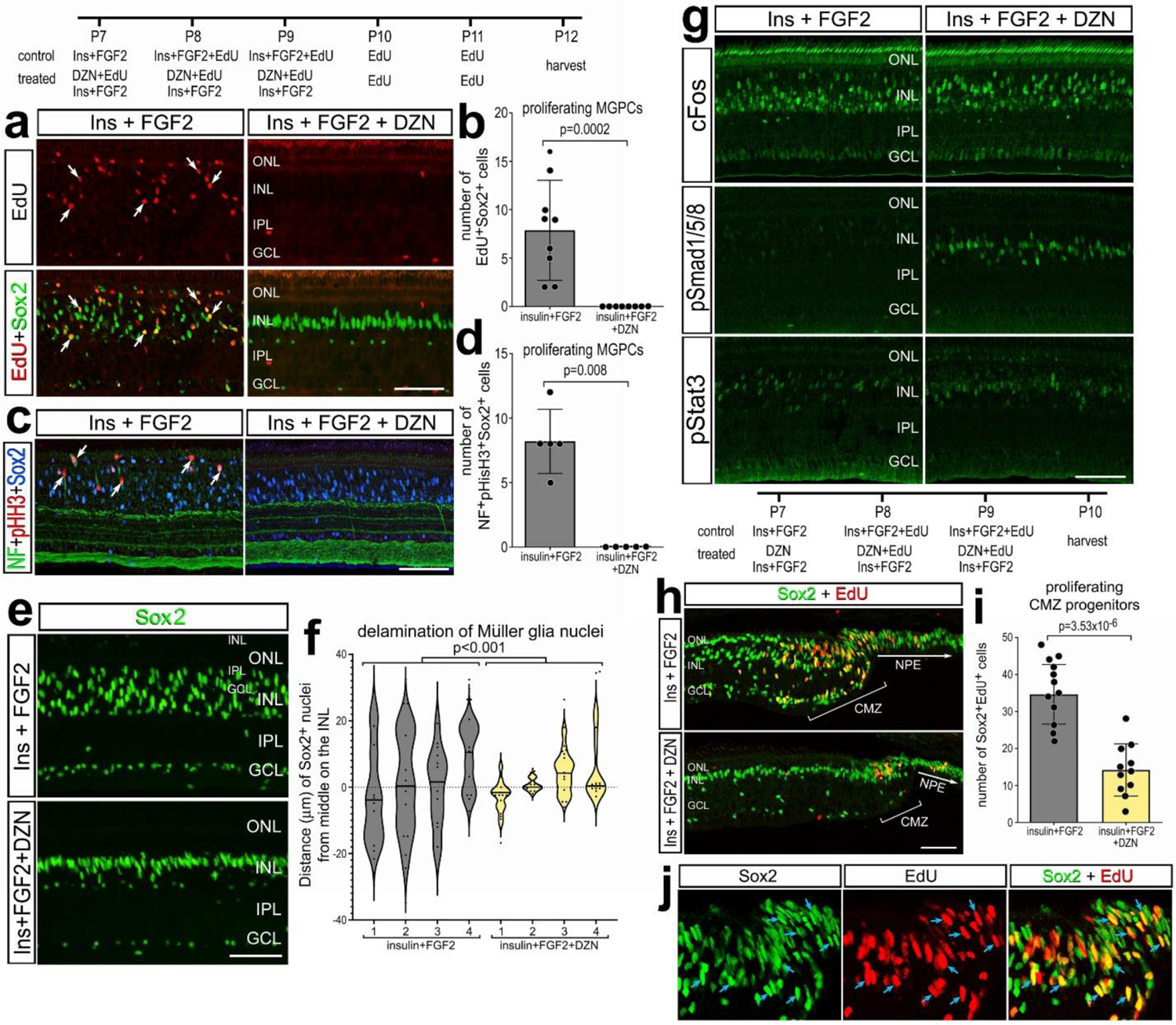
SAHH inhibition in retinas treated with insulin + FGF2 suppresses the proliferation of MGPCs and CMZ progenitors. Eyes were injected with insulin+FGF2+EdU ± DZN at P7, P8 and P9, EdU at P10 and P11, and retinas harvested 24 hrs after the last injection. Sections of the retina were labeled for EdU (red; **b, i, k**), or antibodies to Sox2 (green; **a, e, h, j**; blue; **c**), pHisH3 (red; **c**), cFos (green; **g**), pSmad1/5/8 (green; **g**), or pStat3 (green; **g**). Sections of the far peripheral retina and CMZ were labeled for EdU (red) and Sox2 (green). White arrows indicate the nuclei of MG and blue arrows indicate nuclei in the CMZ that are double-labeled for EdU and Sox2. The calibration bars represent 50 µm. (**b**, **d, i**) Histograms represent the mean (±SD) and each dot represents one biological replicate. The violin histogram in **f** represents a frequency distribution with each dot representing the position of a MG nucleus and line representing the mean for different individual retinas for each treatment. Significance of difference was determined using a Mann-Whitney U test (**b, d**), ANOVA (**f**), or paired t-test (**i**). Abbreviations: NPE – non-pigmented epithelium, CMZ – circumferential marginal zone, ONL – outer nuclear layer, INL – inner nuclear layer, IPL – inner plexiform layer, GCL – ganglion cell layer.

We next investigated whether the proliferation of progenitors in the circumferential marginal zone (CMZ) was influenced by inhibition of SAHH activity by DZN. CMZ progenitors are found at the peripheral edge of the post-hatch chick retina and normally proliferate at relatively low levels, but can be stimulated to proliferate at elevated levels in response to insulin/FGF2 or IGF1 (Fischer and Reh, 2000; Fischer et al., 2002a; Fischer et al., 2008b). We found that DZN-treatment significantly suppressed the proliferation of CMZ progenitors that is stimulated by insulin+FGF2 (Fig. 9i-k). This finding suggests that the activity of HMTs is required for the proliferation of CMZ progenitors, retinal progenitor cells that are distinctly different from MGPCs.

### Inhibition of HMTs in the mouse retina

MG in the adult mouse retina can be reprogrammed to generate neuron-like cells by forced expression of Ascl1 combined with NMDA-induced neuronal damage and HDAC inhibitor to increase chromatin access (Jorstad et al., 2017). Thus, we hypothesized that inhibition of HMTs may influence neurogenic reprogramming in the mouse retina. To test this hypothesis we applied DZN to the retinas of *Glast-CreER:LNL-tTA:tetO-mAscl1-ires-GFP* mice (Jorstad et al., 2017). To induce neuron regeneration from *Ascl1*-overexpressing MG retinas must be damaged by NMDA, and HDAC inhibitor, trichostatin A (TSA), must be applied (Jorstad et al., 2017). Similar to HMT inhibitor, HDAC inhibitor is expected to increase chromatin access. Thus, we replaced TSA with DZN in the treatment paradigm, and tested whether inhibition of HMTs influenced the formation of neurons derived from *Ascl1*-overexpressing MG. *Ascl1-*expression in MG was activated by IP delivery of 4 consecutive daily doses of tamoxifen in adult mice (P80-P90). This was followed by intravitreal injection of NMDA or NMDA+DZN on D8 and injection of vehicle or DZN on D10. Inhibition of HDACs, JAK/STAT, NFkB, TGFB/Smad or ID transcription factors shortly after retinal damaged and forced expression of Ascl1 followed by at least 2 weeks to permit differentiation has been shown to significantly increase differentiation of neuronal cells from MG in adult mice (Jorstad et al., 2017; Jorstad et al., 2020; Palazzo et al., 2022). Accordingly, eyes were harvested and retinas were processed for immunolabeling 2 weeks after the final injection. Inhibition of HMTs in Ascl1-overexpressing MG resulted in no significant change in the de-differentiation of Sox2+ MG (Figs. 10a,c), or differentiation of Otx2+ neuron-like cells (Figs. 10b,d).

**Figure 10.**
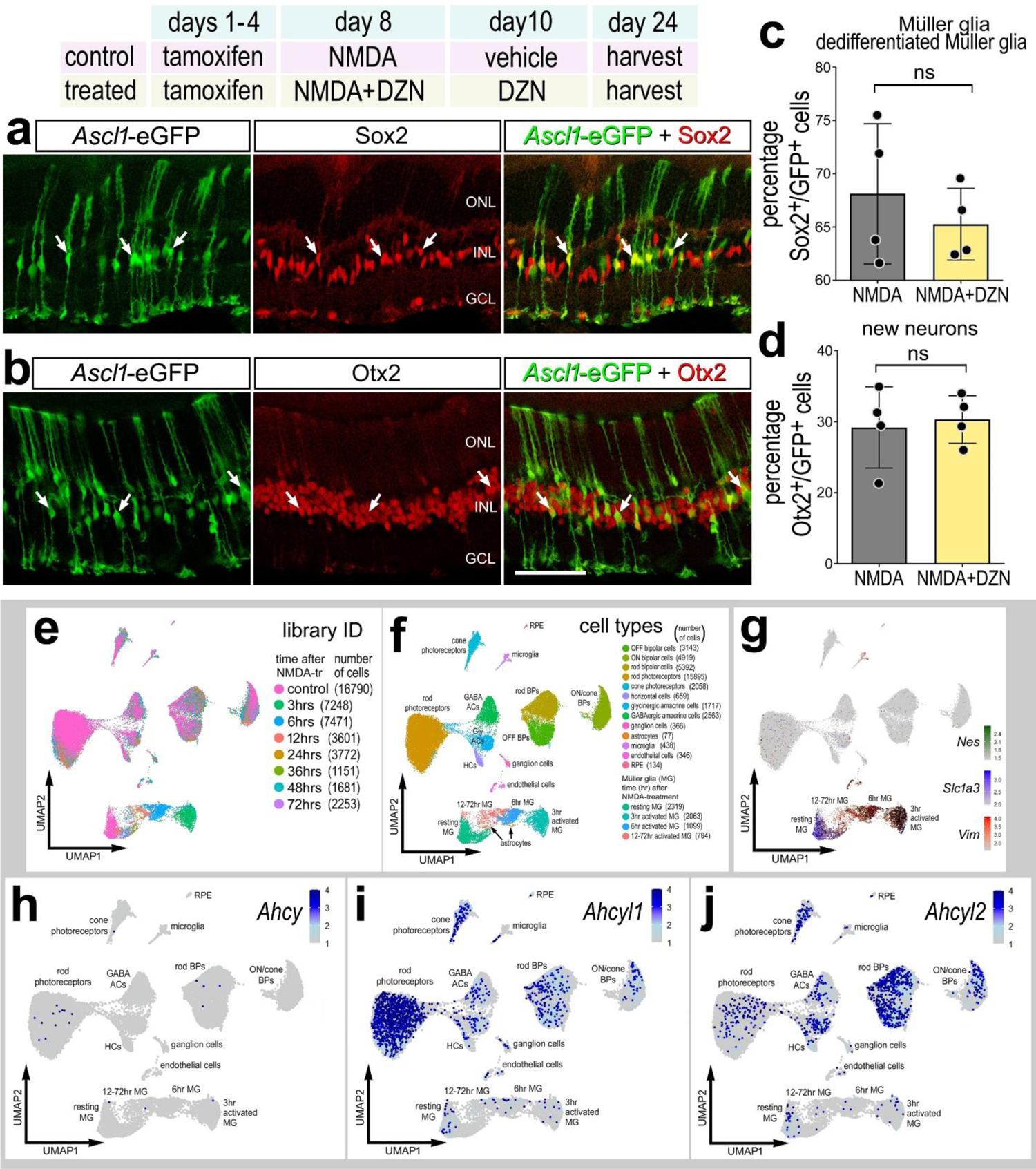
SAHH inhibitor has no effect upon the differentiation of neurons from *Ascl1*-overexpressing MG in the mouse retina. (**a-d**) Tamoxifen was administered IP 1x daily for 4 consecutive days to *Glast*-*CreER:LNL-tTA:tetO-mAscl1-ires-GFP* mice. NMDA was injected intravitreally in left (control) eyes and NMDA + DZN in right (treated) eyes on D8, vehicle ± DZN on D9, vehicle ± DZN on D10, and retinas were harvested 2 weeks after the last injection. Retinal sections were labeled for GFP (green), and Sox2 (red; **a**) or Otx2 (red; **b**). Arrows in **a** and **b** indicate cells double-labeled for GFP and Sox2 or Otx2. Histograms illustrate the mean percentage (±SD and individual data points) of GFP^+^ cells that are Sox2^+^ (**c**) or Otx2^+^ (**d**). Significance of difference (p-values shown) was determined by using a paired t-test. The calibration bar in **b** represents 50 µm. Abbreviations: ONL, outer nuclear layer; INL, inner nuclear layer; GCL, ganglion cell layer. ***Ahcy, Ahcyl1* and *Ahcyl2* in normal and damaged mouse retinas (e-j).** UMAP plots of aggregated scRNA-seq libraries prepared from control retinas and retinas 3, 6, 12, 24, 36, 48, and 72 hours after NMDA damage; numbers of cells per library or per cluster in parentheses (**e**). Clusters of different types of retinal cells were identified based on collective expression of different cell-distinguishing markers as described in the Materials and Methods (**f**). Resting MG and reactive MG identified by expression of *Slc1a3* or *Nes and Vim,* respectively (**g**). UMAP heatmap plots illustrate patterns and expression levels of *Ahcy, Ahcyl1* and *Ahcyl2* across all retinal cells (**h-j**).

To identify expression patterns of *Ahcy*, *Ahcyl1* and *Ahcy2*, we probed scRNA-seq libraries from normal and NMDA-damaged mouse retinas, as described previously (Campbell et al., 2021b; Hoang et al., 2020; Todd et al., 2019). UMAP ordering of cells reveal distinct clusters of different types of retinal cells, with only the MG (*Nes*-, *Slc1a3*- and *Vim*-expressing cells) and distinctly clustered according to time after NMDA-treatment (Figs. 10e-g). UMAP heatmap plots revealed that *Ahcy* is not widely express by any cell types in the mouse retina (Figs. 101h). By comparison, *Ahcyl1* was highly expressed by rod/cone photoreceptors and many inner retinal neurons (Fig. 10i) and Ahcyl2 was highly expressed by many cone photoreceptors and many inner retinal neurons (Fig. 10j). *Ahcy*, *Ahcyl1* and *Ahcy2* were not highly expressed by many resting or activated MG (Fig. 10g-j). Collectively, these findings indicate that DZN-treatment and inhibition of SAHH has no effect upon the reprogramming of Ascl1-overexpressing MG in damaged mouse retinas, likely because *Ahcy, Ahcyl1* and *Ahcyl2* are not high expressed by resting or activated MG.

## Discussion

We find that *AHCY, AHCYL1, AHCYL2* and HMTs are dynamically expressed by MG in damaged retinas. Dynamic changes of mRNA levels are strongly correlated with changes in protein levels and function (Liu et al., 2016). In the chick retina MG upregulated downregulate *AHCY* genes during the transition to MGPCs. By contrast, *AHCY*s were not widely expressed by MG in damaged mouse retinas. Our findings indicate that inhibition of SAHH effectively reduced levels of histone methylation and potently suppress the formation of proliferating MGPCs in the chick retina (Fig. 11). By comparison, inhibition of SAHH had no effect upon the differentiation of neuron-like cells from Ascl1-over expressing MG in damaged mouse retinas. In the chick, inhibition of SAHH had significant effects upon gene expression and chromatin access in MG and microglia (Fig. 11). Many changes in expression and access are associated with modules of genes that influence retinal development, neuronal differentiation and cell signaling pathways that are known to influence the formation of proliferating MGPCs. (Du et al., 2021; Kong et al., 2014). In summary, the effects of SAHH inhibitor are consistent with the notion that the activity of HMTs and changes in chromatin access are required for the formation of proliferating MGPCs in the chick retina.

**Figure 11.**
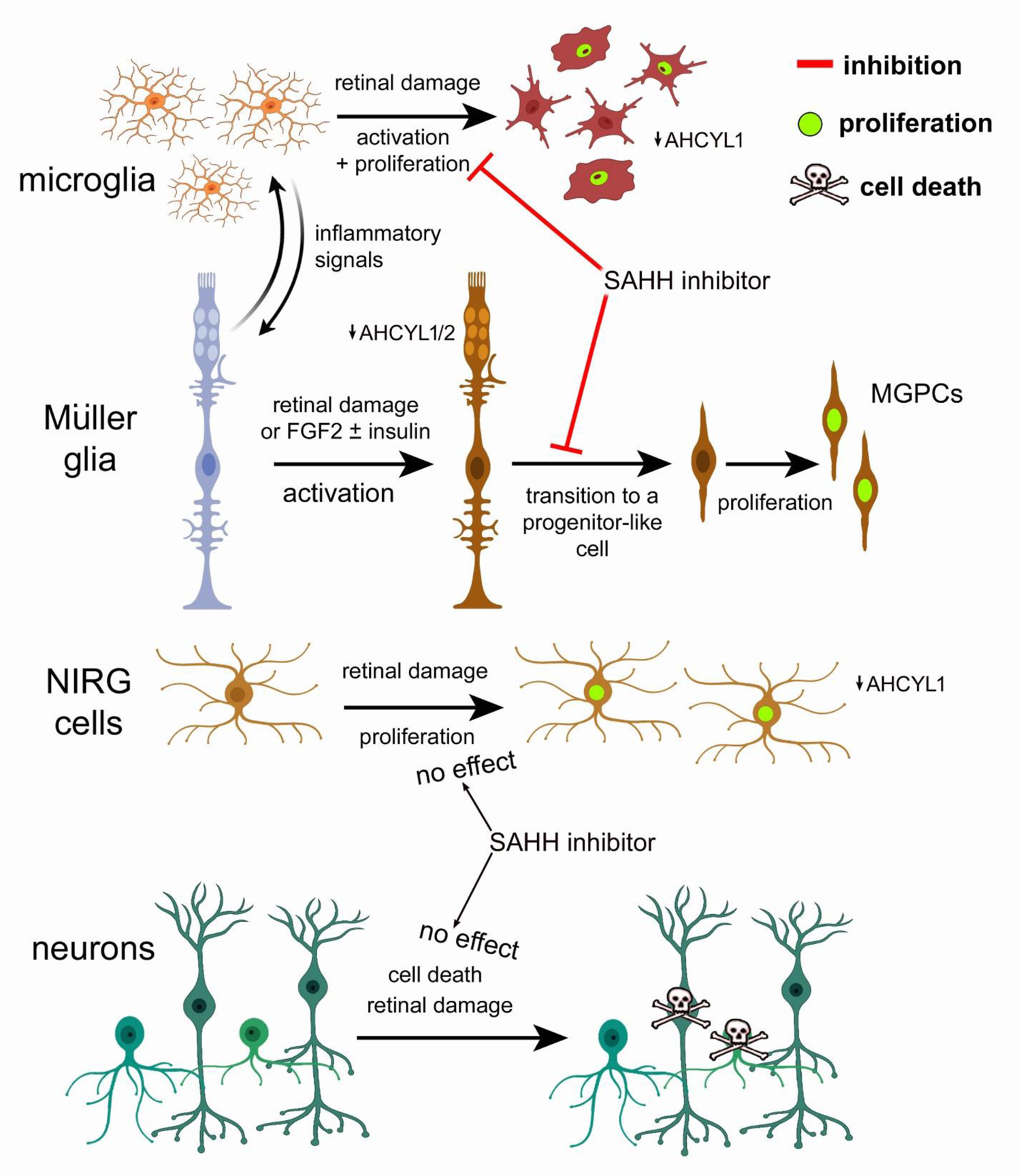
Schematic summary of actions of SAHH inhibitor on microglia, MG, NIRG cells and neurons. This figure was created with cells from <u>BioRender.com</u>.

### NMDA-induced retinal damage influences chromatin accessibility in MG

For scATAC-seq libraries, UMAP ordering of cells from control and damaged retinas indicated that MG, unlike other types of retinal cells, had significant differences in chromatin access represented by spatial separation of MG from different treatment groups. We identified more than 480 genes with differential chromatin access in MG from NMDA-damaged retinas. These genes included decreased access to many different immediate early genes and genes associated with morphogenesis and neurogenesis, and increased access to genes associated with cell signaling pathways, such as TGFβ, gp130/Jak/Stat and Notch, which are known to influence glial reactivity and formation of MGPCs (Ghai et al., 2010; Todd et al., 2016a; Todd et al., 2017). Changes in chromatin access in glial cells have been well-described during development. For example, the maturation of astrocytes involves extensive changes in chromatin access (Lattke et al., 2021). Similarly, developmental differences in gene expression and chromatin access have been identified and compared across retinal progenitors and MG, and prevalent changes in chromatin access demonstrated for neurogenic genes (VandenBosch et al., 2020).

### Genes affected by inhibition of SAHH

We observed a modest correlation of genes with increased/decreased chromatin access and increased/decreased expression levels. There were few genes with increased chromatin access and downregulated gene expression. It is possible that heterogeneity of cells within UMAP-clustered MG accounts for diminished correlation between genes with chromatin access and expression levels. A combined multi-omic approach may resolve this possibility wherein individual cells receive unique DNA barcodes and unique molecular identifiers to measure both chromatin access and gene expression levels. By comparison, we observed a strong correlation between chromatin access, mRNA levels, and motif access in MG for transcription factors related to glial identity and retinal development. Similarly, a recent stud*y* identified genomic regions important to corticogenesis by assessing the activity of gene-regulatory elements by mapping, at single-cell resolution, gene expression and chromatin access (Trevino et al., 2021). Networks of gene regulation, including expression programs of glial lineages, were strongly correlated between gene-regulatory elements and expression levels for key transcription factors (Trevino et al., 2021).

### Inhibition of SAHH and proliferation of progenitors

SAHH inhibitor potently suppressed the formation of proliferating MGPCs in damaged retinas, suppressed the formation of MGPCs in retinas treated with insulin and FGF2 in the absence of damage, and suppressed the proliferation of stimulated CMZ progenitors at the far peripheral edge of the retina. Collectively, these findings suggest that the activity of SAHH and HMTs are required for the proliferation of retinal progenitor cells. Consistent with labeling for proliferation markers in tissue sections, SAHH inhibition significantly downregulated many proliferation-related genes in MG in damaged retinas, but did not significantly influence chromatin access to proliferation-related genes. These observations suggest the changes in chromatin access and gene expression caused by inhibiting SAHH act upon genes that are upstream of regulating re-entry into the cell cycle. However, it is possible that inhibition of SAHH affected methyltransferases in addition to HMTs to suppress the formation of MGPCs.

### Inhibition of SAHH and cell signaling in MG

Inhibition of SAHH in undamaged retinas stimulated MG to upregulate genes associated with resting MG, pro-glial transcription factors, and cell signaling pathways known to stimulate and suppress MGPC formation. For example, treatment with SAHH inhibitor resulted in upregulation of some of the most highly expressed genes in resting MG including *GLUL, RLBP1, CSPG5* and *GPR37*. In addition, treatment with SAHH inhibitor upregulated transcription factors, such as *ID4* and *NFIA/C/X*, which are known to promote glial differentiation and maintain glial identity (Clark et al., 2019; Deneen et al., 2006) and transcription factors, such as *SIX3* and *SIX6*, which are known to maintain retinal progenitor cells (Diacou et al., 2018). Further, we observed upregulation of Wnt suppressors such as *WIF1*, *SFRP1* and *SFRP2,* which are expected to suppress MGPC formation, since Wnt-signaling promotes the proliferation of MGPCs (Gallina et al., 2016). However, the SAHH inhibitor did not induce the formation of MGPCs in undamaged retinas, suggesting that the upregulation of pro-glial networks supersedes any upregulation of pro-MGPC cell signaling pathways. Collectively, these findings indicate inhibition of SAHH in undamaged retinas predominantly activates networks that promote glial phenotype and reprogramming into progenitors.

Inhibition of SAHH in undamaged retinas caused MG to downregulate genes associated with cell signaling pathways expected to promote MGPC formation, including midkine/pleotrophin (*MDK, PTN, SDC4*) (Campbell et al., 2021b), Jak/Stat-signaling (*SOCS3*) (Todd et al., 2016a), Notch-signaling (*MAML2*) (Ghai et al., 2010; Hayes et al., 2007) and MMPs (*TIMP2*) (Campbell et al., 2019). DZN-treatment of MG in NMDA-damaged retinas resulted in upregulation of transcription factor *HES1* and downregulation of *ID3,* which are known to promote glial development and identity (Furukawa et al., 2000; Wall et al., 2009). By comparison, *MSX1* is downregulated in MG with SAHH inhibition; MSX1 is expected to promote the phenotype of peripheral retinal progenitors (Belanger et al., 2017). Nevertheless, SAHH inhibition in undamaged retinas did not result in the formation of MGPCs. Although inhibition of SAHH stimulated MG in undamaged retinas to upregulate gene modules associated with neurogenesis and pro-proliferation, the context is changed in damaged retinas and some of these effects are absent or opposite. For example, gene modules associated with cell migration, PI3K-Akt-signaling and cell cycle are upregulated in MG in undamaged retinas whereas these gene modules are downregulated in MG in damaged retinas. We found that SAHH inhibition influenced gene expression and chromatin accessibility in undamaged retinas, wherein levels of *AHCYL1* and *AHCYL2* were relatively high. These effects of SAHH inhibitor on MG may result from: (i) inhibition of HMT activity in MG, (ii) inhibition of the methlytransferases in addition to HMTs, (iii) off-target effects of DZN, or (iv) indirect effects of SAAH inhibition on microglia or retinal neurons. DZN is known to inhibit S-adenosylhomocysteine hydrolase (SAHH) (reviewed in (Vizán et al., 2021), but is widely described in the literature as an EZH2 inhibitor. *AHCY* is widely expressed by retinal neurons and MG; levels of expression are high in resting glia, downregulated in activated glia, and unchanged in retinal neurons following damage (not shown). SAHH regulates the concentration of intracellular levels of S-adenosylmethionine which is required for transmethylation reactions and levels of SAH which inhibits transmethylation reactions, including reactions mediated by HMTs (Vizán et al., 2021).

Inhibition of SAHH in damaged retinas resulted in increased autocrine LR-interactions in MG for pathways involving SFRP1 (to inhibit Wnt) and Jagged-Notch. Wnt-signaling stimulates the proliferation of MGPCs in zebrafish (Meyers et al., 2012; Ramachandran et al., 2011), chick (Gallina et al., 2016) and mouse retinas (Osakada et al., 2007; Yao et al., 2016). The relationship of Notch-signaling to the formation of proliferating MGCPs is more complicated and appears to require dynamic regulation. In short, evidence suggests that Notch-signaling maintains glial phenotype in resting glia, downregulation is required early for de-differentiation, upregulation is required for proliferation of MGPCs and downregulation is required for neuronal differentiation (Campbell et al., 2021a; Campbell et al., 2022b; Ghai et al., 2010; Hayes et al., 2007; Lee et al., 2020; Sahu et al., 2021). Many of the autocrine LR-interactions induced by SAHH inhibitor in damaged retina, including those related FGFR1, Notch and midkine-signaling, are expected to promote the formation of MGPCs (Campbell et al., 2021b; Fischer et al., 2009a; Ghai et al., 2010). Similarly, we observed upregulation of many genes associated with different cell signaling pathways, such as *SPRY2* and *IGFBP5,* which are expected to suppress the formation of MGPCs by inhibiting MAPK signaling (Fischer et al., 2009a; Fischer et al., 2009b). By comparison, we observed downregulation of *BMP4* and *SMAD5* which is consistent with suppressed formation of MGPCs (Todd et al., 2017). In summary, DZN-induced changes in genes and modules associated with cell signaling pathways known to influence the formation of MGPCs are not entirely consistent with suppressing the formation of MGCPs.

Although changes in gene expression and putative LR-interactions implicated changes in cell signaling pathways, we did not observe decreased activation of cFos, Jak/Stat- or BMP/Smad-signaling in MG from damaged retinas treated with SAHH inhibitor. However, we did observed a significant decrease in mTor-signaling in MG in damaged retinas treated with SAHH inhibitor; wherein mTor-signaling is known to be required for the formation of MGPCs in damaged chick retinas (Zelinka et al., 2016). By comparison, inhibition of Fatty acid-binding proteins (FABPs) or Fatty acid synthase (FASN) potently reduces levels of pS6, pStat3 and pSmad1/5/8 in MG in damaged retinas (Campbell et al., 2022a). Collectively, these findings suggest the SAHH inhibition predominantly acts by modulating chromatin access, changes in expression of pro-glial transcription factors, and access to TF motifs, rather than influencing cell signaling.

### Inhibition of SAHH in damaged mouse retinas

Changes in chromatin access and histone methylation mediated by EZH2 in the developing mouse retina are required for the formation of MG (Iida et al., 2015; Zhang et al., 2015, 2). Inhibition of SAHH is expected to block the activity of EZH2. However, SAHH inhibitor failed to influence the reprogramming of Ascl1-overexpressing MG in the mouse retina. However, SAHH and EZH2 (not shown) are not widely expressed by resting or activated MG in the mouse retina. Thus, it is not surprising that the SAHH inhibitor had no effect on the formation of neuron-like cells from Ascl1-overexpressing MG in damaged mouse retinas.

## Conclusions

Our findings indicate that the activity of SAAH is necessary for the formation of proliferating MGPCs and the proliferation of CMZ progenitors in the chick retina. Inhibition of SAAH influenced gene networks through changes in chromatin access and gene expression, and these changes were manifested in MG and microglia. Activation of cell signaling pathways known to promote the formation of MGPCs were largely unaffected by SAHH inhibition, with the exception of signaling through mTor. Collectively, our findings indicate that inhibition of SAHH results in changes in expression and chromatin access for genes that are known to promote or inhibit gliogenesis and MGPC formation. Increases in expression/access to genes that inhibit MGPC formation supersede those that promote this process. By comparison, inhibition of SAHH had no effect upon the formation of neuron-like cells from Ascl1-overexpressing MG. We propose that the activity of SAAH and HMTs may be among the key differences in avian and mammalian MG that is required for chromatin remodeling that promotes the formation of MGPCs and retinal regeneration.

## Acknowledgements

This work was supported by RO1 EY022030-10 (AJF). We thank Drs. Tim Stuart and Jared Tangeman for providing assistance with establishing appropriate reference genomes for scATAC-seq libraries. We thank Dr. Tom Reh for kindly providing transgenic mice. We thank Olivia Taylor for comments that helped to shape the final from of the paper.

## Author contributions

WAC and HME – experimental design, execution of experiments, collection of data, data analysis, construction of figures, bioinformatics, preparation of RNA-seq and ATAC-seq and writing the manuscript. DT, ECH, LEK, LV and DA – execution of experiments and collection of data. AJF – experimental design, data analysis, bioinformatics, construction of figures and writing the manuscript.

## Data availability

Gene-Cell matrices for scRNA-seq data for libraries from saline and NMDA-treated retinas are available through GitHub: https://github.com/jie[wwwang/Singlecell-retinal-regeneration](http://www.ang/Singlecell-retinal-regeneration)

Gene-Cell matrices for scRNA-seq data for libraries for NMDA-treated retinas and DZNexp4-treated retinas are available through GitHub: https://github.com/fischerlab3140/ Gene-Cell matrices for scATAC-seq data are available through GitHub: https://github.com/fischerlab3140/

scRNA-Seq data for chick retinas treated with NMDA or insulin and FGF2 can be queried at: https://proteinpaint.stjude.org/F/2019.retina.scRNA.html.

---

**Supplemental Figure 1.**
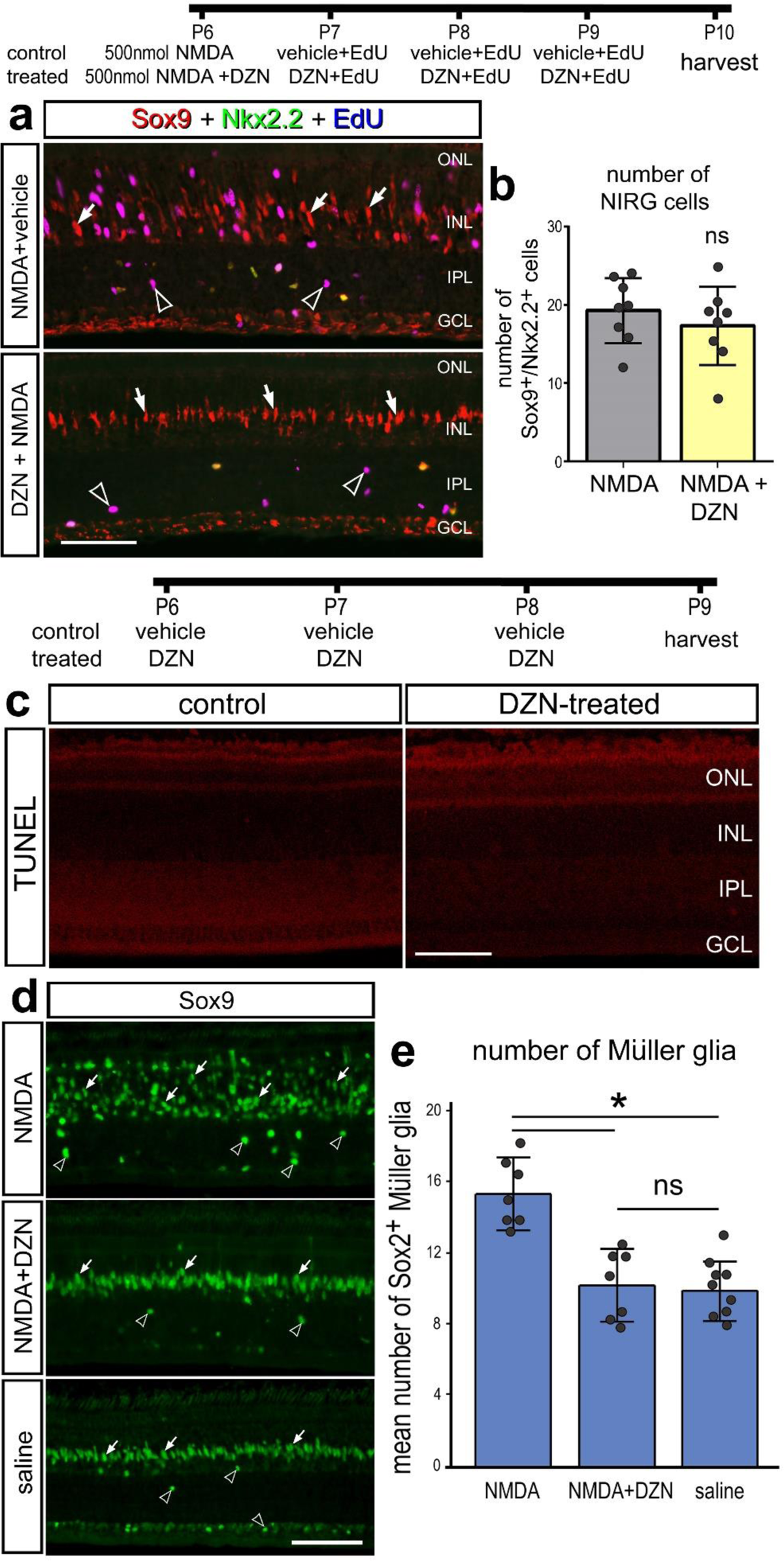
NIRG cells, cell death and numbers of MG are unaffected by EZH2 inhibitor. (**a-b**) Eyes were injected with NMDA ± DZN (EZH2 inhibitor) at P6, EdU ± DZN at P7, P8 and P9, and retinas harvested at P10. Sections of the retina were labeled for EdU (blue - **a**), Nkx2.2 (green – **a**) and Sox9 (red - **a**). (**c-e**) Eyes were injected with vehicle or DZN at P6, P7 and P8, and eyes were harvested at P9. Sections of the retina were labeled for dying cells (TUNEL - **c**) or Sox9 (green – **d**). Histograms represent the mean (±SD) and each dot represents on biological replicate for numbers of NIRG cells (**b**) or numbers of Sox9^+^ MG (**d**). Significance of difference (*p<0.05, ns – not significant) was determined using a paired t-test (**b**) or ANOVA with post-hoc t-test with Bonferroni correction (**e**). Arrows indicate the nuclei of MG and hollow arrowheads indicated the nuclei of NIRG cells (**a,d**). Abbreviations: ONL – outer nuclear layer, INL – inner nuclear layer, IPL – inner plexiform layer, GCL – ganglion cell layer.

